# Processing and Storage Methods Affect Oral and Gut Microbiome Composition

**DOI:** 10.1101/2023.06.13.544865

**Authors:** Dorothy K. Superdock, Wei Zhang, Angela C. Poole

**Author notes:** Address correspondences to Angela C. Poole.

## Abstract

Across microbiome studies, fecal and oral samples are stored and processed in different ways, which could affect the observed microbiome composition. Here, we compared treatment methods, which included both storage conditions and processing methods, applied to samples prior to DNA extraction to determine how each affects microbial community diversity as assessed by 16S rRNA gene sequencing. We collected dental swab, saliva, and fecal samples from 10 individuals, with three technical replicates per treatment method. We assessed four methods of processing fecal samples prior to DNA extraction. We also compared different fractions of frozen saliva and dental samples to fresh samples. We found that lyophilized fecal samples, fresh whole saliva samples, and the supernatant fraction of thawed dental samples retained the highest levels of alpha diversity in samples. The supernatant fraction of thawed saliva samples had the second highest alpha diversity compared to fresh. Then we investigated the differences in microbes between different treatments at the domain and phylum levels as well as identified the amplicon sequence variants (ASVs) that were significantly different between the methods producing the highest alpha diversity and the other treatment methods. Lyophilized fecal samples had a greater prevalence of Archaea as well as a greater ratio of Firmicutes to Bacteroidetes compared to the other treatment methods. Our results provide practical considerations, not only for selection of processing method, but also for comparing results across studies that use these methods. Our findings also indicate differences in treatment method could be a confounding factor influencing the presence, absence, or differential abundance of microbes reported in conflicting studies.

## INTRODUCTION

Assessment of oral and gut microbial communities has become prevalent in clinical studies as therapeutics targeting the microbiome are increasingly explored (Wade, 2013; Lynch and Pedersen, 2016). Methods used during the workflow while generating microbial genetic sequencing data can vary widely between studies, including sample collection method, storage, and DNA extraction, which can all influence observed microbial composition (Hugerth and Andersson, 2017; Kim et al., 2017; Vandeputte et al., 2017; Teng et al., 2018; Fiedorová et al., 2019; Cunningham et al., 2020; Marotz et al., 2021). If these methods affect detection of the microbes that mediate the disease or treatment being studied, research groups may report conflicting results. It is therefore important to consider the impact of differences in storage and processing methods before making comparisons across studies. The objective of this current study was to compare microbiome composition in a human subjects cohort across different fecal processing methods as well as different saliva and dental sample fractions present after thawing from −20°C to determine which methods maximized observed microbial diversity and compare how the observed microbiomes differed between methods.

We collected samples from 10 individuals and each sample was subjected to several different treatments, including both storage and processing conditions (Figure 1), in triplicate, prior to DNA extraction and 16S rRNA gene sequencing. For fecal samples, we investigated how lyophilization differed from other treatments because this has become a more common long-term preservation method. Due to the removal of water, lyophilization both reduces sample mass and allows for indefinite storage at room temperature without further microbial growth. We compared lyophilized fecal samples to: 1) fresh fecal samples (from which DNA was extracted on the day of collection without first being frozen), 2) fecal samples frozen at −80°C then subsequently ground in liquid nitrogen (LN2), which is a common method to homogenize a frozen fecal sample to a fine powder to prepare it for DNA extraction, as well as 3) fecal samples that were left out at room temperature prior to freezing at −80°C and then ground in liquid nitrogen (LN2post72hr), simulating studies during which human participants ship their fecal samples to investigators (Song et al., 2016), who then freeze the samples until further processing. In this study, we used a 72 hour incubation time, which is three days at room temperature.

**Figure 1.**
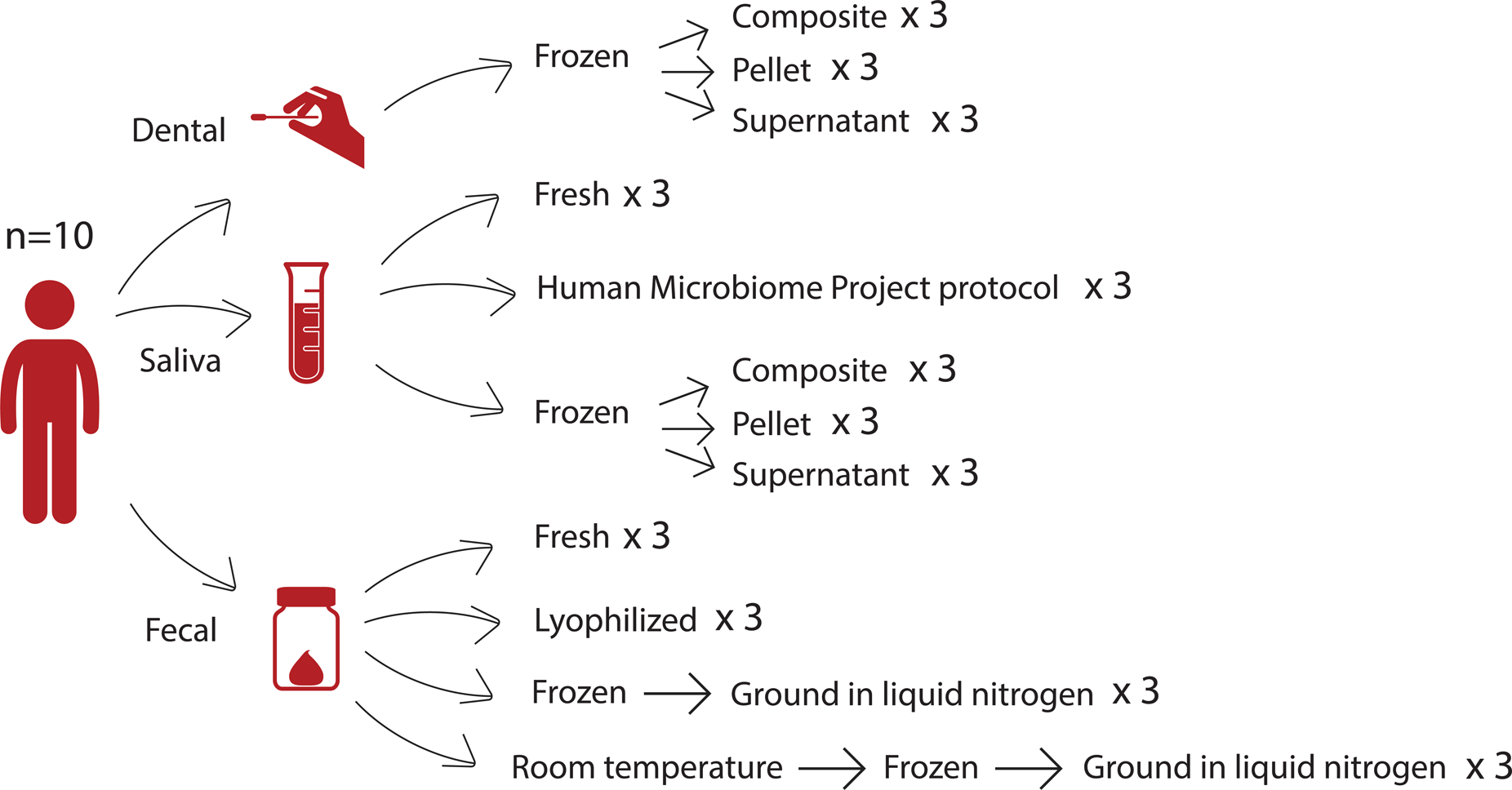
Study design. From 10 donors, we collected three different sample types—dental swabs, saliva samples, and fecal samples—which were then divided into several downstream treatment methods. Treatments included different storage types, sample fractions, and processing methods. For each sample, each treatment was tested in triplicate.

We also examined differences between thawed fractions of oral samples, specifically saliva samples and dental swabs. Saliva samples are commonly collected to determine overall oral microbiota composition. For studies involving caries or periodontal disease, dental swabs can be collected to enhance detection of bacteria that form biofilms on teeth in a less invasive manner than collecting subgingival plaque (Lu et al., 2022; Uyghurturk et al., 2022; Selway et al., 2023).

These sample types are commonly frozen then thawed before processing. Upon thawing, there is often precipitation of proteins, such as mucins, present in the samples (Schneyer, 1956). Researchers may make different decisions at the input step for a DNA extraction protocol regarding whether to use centrifugation to separate the sample into a pellet and supernatant or vortexing to homogenize the sample. We compared the three different fractions of thawed saliva and dental samples (pellet, supernatant, or composite) to determine which part of the sample should be targeted for DNA extraction if the goal is to maximize alpha diversity. The supernatant and pellet fractions resulted from centrifugation of the sample, and the composite fraction was a vortexed mixture of the whole sample prior to the centrifugation step used to separate the supernatant and pellet fractions. In addition, for saliva samples we evaluated the Human Microbiome Project Phase 1 (HMP) “Initial Processing of Saliva” protocol, which was developed by the HMP as part of a common set of guidelines and protocols (https://www.hmpdacc.org/), as well as a fresh condition where samples were subject to DNA extraction on the day of collection without first being frozen.

## MATERIALS AND METHODS

### Human Subjects

We collected fecal, saliva, and dental swabs from each of ten healthy participants (n=6 females, n=4 males) between 25-50 years of age (Cornell University IRB protocol #1106002281). Samples were treated using several different processing and storage methods as shown in Figure 1 and described below. Hereafter we refer to these different combinations of processing and storage methods as treatments.

### Saliva samples

Each participant expressed approximately 5 mL of saliva into a 50 mL conical. Each saliva sample was vortexed and 1) 500-750 μl was pipetted into a Powersoil Bead Tube (MOBIO PowerSoil DNA Isolation Kit, QIAGEN Cat #12888) for DNA extraction within two hours of collection (fresh), 2) 1 mL was pipetted into a microcentrifuge tube and was centrifuged at 2600 x g for 15 minutes at room temperature, then 750 μL of supernatant was added to 750 μL of Powersoil Bead Buffer and frozen at −80°C prior to DNA extraction (HMP), and 3) 1 mL was pipetted into another microcentrifuge tube and was frozen without centrifugation. After thawing frozen aliquots, samples contained varying amounts of precipitate and were vortexed so that an aliquot containing both liquid and precipitate was removed with a wide orifice pipette tip to represent the composite fraction. Then samples were centrifuged at 1500 x g at 4°C for 15 minutes to separate a pellet from the supernatant fraction.

### Dental swabs

Participants were asked to simulate brushing the buccal side of their teeth, top and bottom rows, for 30 seconds with nylon swabs (Epicentre, cat # QEC89100). Six dental swabs were collected per participant. Dental swabs were placed in Bead Solution from the MOBIO PowerSoil kit for 5-10 minutes, swirled, and vortexed briefly on the lowest setting in order to facilitate the transfer of dental swab material into the bead solution. 100-200 uL of Bead Solution containing dental swab material was then combined for each participant, split into triplicate, then frozen at −80°C within 90 minutes of collection. Upon thawing, each of the three samples was first mixed to obtain the composite aliquot then centrifuged at 1500 x g at 4°C for 15 minutes to separate a pellet from the supernatant fraction.

### Stool samples

Participants collected their own stool samples using a commode specimen collection kit (ThermoFisher, cat # 02-544-208). After receipt by the lab on the day of collection, each stool sample was mechanically homogenized in a zip-top freezer bag using a rolling pin, and transferred into 50 mL conical tubes containing no preservative or buffer for further processing downstream using four different methods. Samples were 1) fresh, 2) lyophilized (freeze dried), 3) frozen at −80°C for > 2 weeks, then ground in liquid nitrogen (LN2), or 4) stored at room temperature (RT) for 72 hours, frozen at −80°C for > 2 weeks, then finally ground in liquid nitrogen (LN2post72hr).

### 16S rRNA library generation and sequencing

Microbial DNA was extracted from all gut and oral samples using the MO BIO PowerSoil-htp Soil DNA Isolation Kit (MO BIO Laboratories, cat # 12955-4) according to the manufacturer’s protocol except instead of vortexing, samples were placed in a BioSpec 1001 Mini-Beadbeater-96 for two minutes after the addition of Solution C1. For amplification of the V4 region of the 16S rRNA gene we used 10-50 ng of sample DNA in duplicate 50 μl PCR reactions with 5 PRIME HotMasterMix and 0.1 μM forward (515F) and reverse (806R) primers using the PCR program previously described (Caporaso et al., 2011) but with 25 cycles. We purified amplicons using the Mag-Bind E-Z Pure Kit (Omega Bio-tek, cat # M1380) and quantified with Invitrogen Quant-iT PicoGreen dsDNA Reagent. 100 ng of amplicons from each sample were pooled and sequenced on an Illumina MiSeq instrument using 2×250 bp paired end sequencing. We used QIIME2 version 2022.2 (Bolyen et al., 2019) to perform microbiome bioinformatics. We demultiplexed and quality filtered raw sequence data using q2-demux, denoised using DADA2 (Callahan et al., 2016) via q2-dada2, created a phylogenetic tree using q2-fragment-insertion, and calculated diversity metrics using q2-diversity after gut samples were rarefied to 31915 sequences per sample and oral samples were rarefied to 19360 sequences per sample. For analyses where saliva samples and dental samples were analyzed separately, diversity metrics were calculated separately. (For microbial diversity analyses, we excluded the HMP condition for saliva samples to avoid unbalanced groups.) To assign taxonomy to ASVs, we used the classify-sklearn naïve Bayes taxonomy classifier via the q2-feature-classifier plugin (Bokulich et al., 2018) against Greengenes 13_8 99% OTUs reference sequences (McDonald et al., 2012). Although samples were all processed in triplicate, some of the sample replicates failed at some point during the library preparation workflow and failed to produce sufficient yields for sequencing. Additionally, only 12 out of the 30 samples in the HMP condition made it to the sequencing step of the library preparation workflow while the rest failed to produce sufficient yields for sequencing. In total, for the rest of the saliva samples, there were 28 fresh, 30 composite, 30 pellet, and 25 supernatant samples included in the analyses. For fecal samples, there were 30 lyophilized, 30 fresh, 30 LN2, and 30 LN2post72hr samples included in the analyses. For dental swabs, there were 23 composite, 28 pellet, and 28 supernatant samples included in the analyses.

### Statistical analysis

We performed all statistical analyses in RStudio using R version 4.2.1 (R Core Team, 2022). To determine how microbiome composition differs at the phylum and domain levels, we used taxa exported from taxa bar plots generated by QIIME2 and used the lme4 R package to fit linear mixed models with treatment as a fixed effect and sample donor as a random effect, e.g. lmer(Archaea Relative Abundance ∼ Treatment + (1|SubjectID), separately for each sample type. For fecal samples, we tested whether the Firmicutes:Bacteroidetes ratio was different depending on processing method by fitting the following model: FBratio ∼ Treatment + (1|SubjectID). We performed PERMANOVA analysis to compare within-group to between-group beta diversity distances using adonis2 of the vegan R package (Oksanen et al., 2022) and pairwise.adonis2 (Martinez Arbizu, 2017), stratifying by donor whenever donor was not used as a fixed effect in the models, i.e. using the parameters “permutations=” (adonis2) or “strata=” (pairwise.adonis2) in the model formulas. Principal coordinates analysis (PCoA) plots were generated using vegan (Oksanen et al., 2022). We performed alpha diversity analyses using the lme4 R package to fit linear mixed models using each alpha diversity metric—Faith’s Phylogenetic Diversity (Faith’s PD, phylogenetic diversity), Pielou’s Evenness (evenness), or Observed ASVs—as the response variable, treatment as a fixed effect, and subject as a random effect, e.g. using the following equation for each alpha diversity metric: Alpha Diversity ∼ Treatment + (1|SubjectID). We performed this analysis separately for each sample type. For all linear mixed models, we performed post-hoc pairwise comparisons employing the emmeans R package using the Tukey method for p-value adjustment. For the alpha diversity correlation analysis, we averaged Faith’s PD for saliva samples and dental samples within each subject and performed a Pearson correlation. For the saliva and fecal sample correlation analysis we selected one sample replicate for fresh saliva samples and one sample replicate for lyophilized fecal samples (based on the highest ASV count), then filtered ASVs to only include those that were present in greater than 50% of participants. We then performed a Pearson correlation between saliva and fecal ASVs in each participant. We FDR-corrected p-values using a Benjamini-Hochberg correction and considered an adjusted p-value (q-value) of <0.05 to be statistically significant. We used MaAsLin2 (Mallick et al., 2021) including all sample replicates per each treatment condition to evaluate the effect of treatment type on the relative abundances of microbes.

## RESULTS

### Microbiome composition differs at the phylum and domain levels between fecal, dental, and saliva treatment methods

Treatment effects were observed at the phylum level. Notably, in the fecal samples, there were differences in the proportions of the top two dominant phyla in the gut, Firmicutes and Bacteroidetes (Figure 2A). Compared to the other three treatments (fresh, LN2, LN2post72hr), lyophilization was associated with a greater than two-fold increase in the ratio of Firmicutes to Bacteroidetes (p<0.0001). Lyophilized samples also had a higher relative abundance of Actinobacteria compared to all other treatments (p<0.0001). LN2post72hr samples had a higher relative abundance of Actinobacteria compared to fresh as well (p=0.03). Fresh samples had a higher relative abundance of Tenericutes compared to lyophilized (p=0.01) and LN2 samples (p=0.03). Fresh samples also had a higher relative abundance of Proteobacteria compared to lyophilized samples (p=0.049). Additionally, there were differences observed at the highest taxonomic rank (domain) level in fecal samples, as detectable by 16S rRNA sequencing. Archaea were only detected in 36 out of 120 total fecal samples and only in five of the ten donors. In two of these five donors, Archaea was detected in only a single replicate sample: 6 out of 133346 ASVs and 18 out of 102727 ASVs. Within the samples from the remaining three donors, lyophilization was associated with an increased relative abundance of Archaea compared to fresh (p=0.009), LN2 (p=0.004), and LN2post72hr (p=0.04).

**Figure 2.**
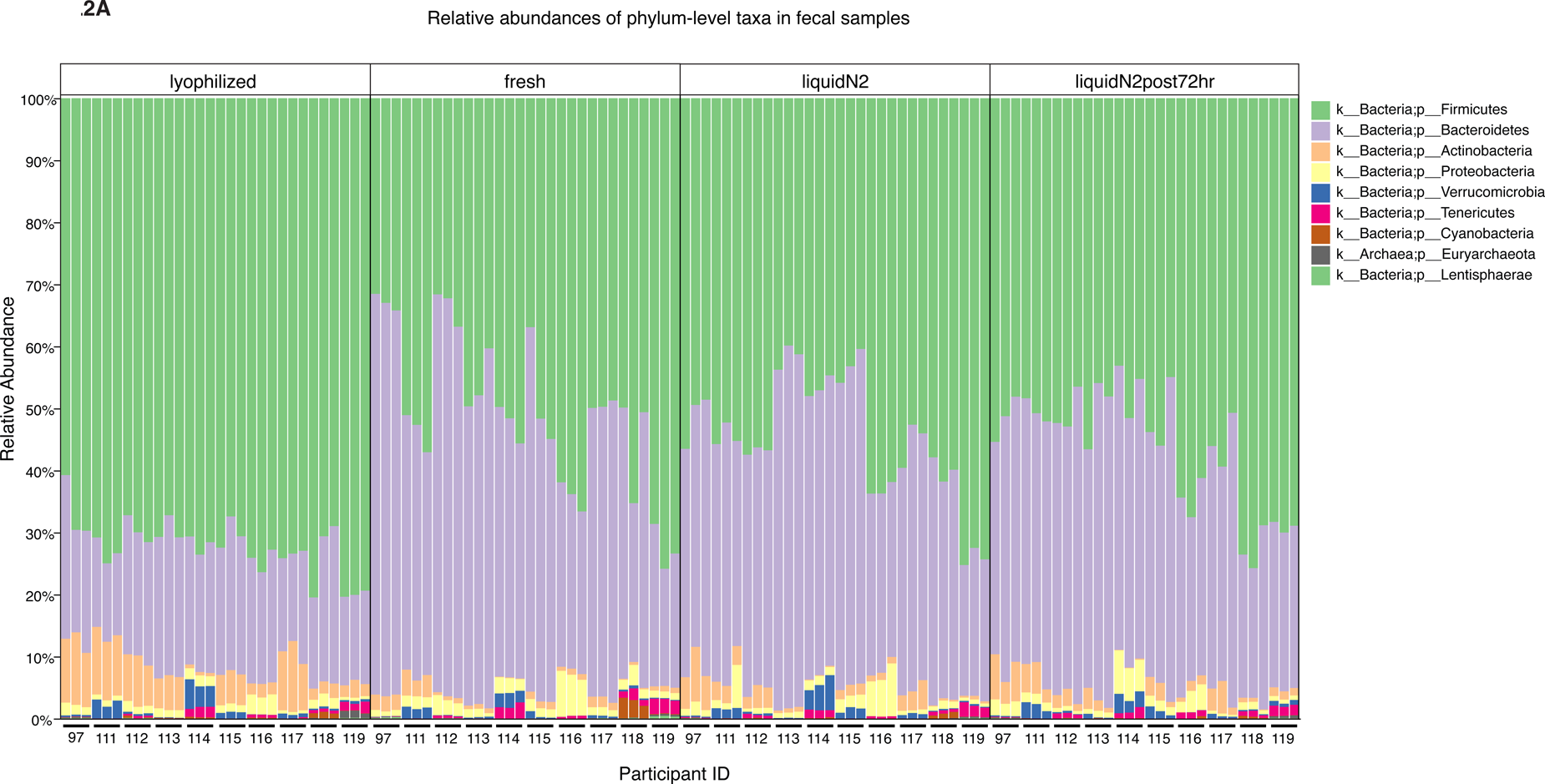
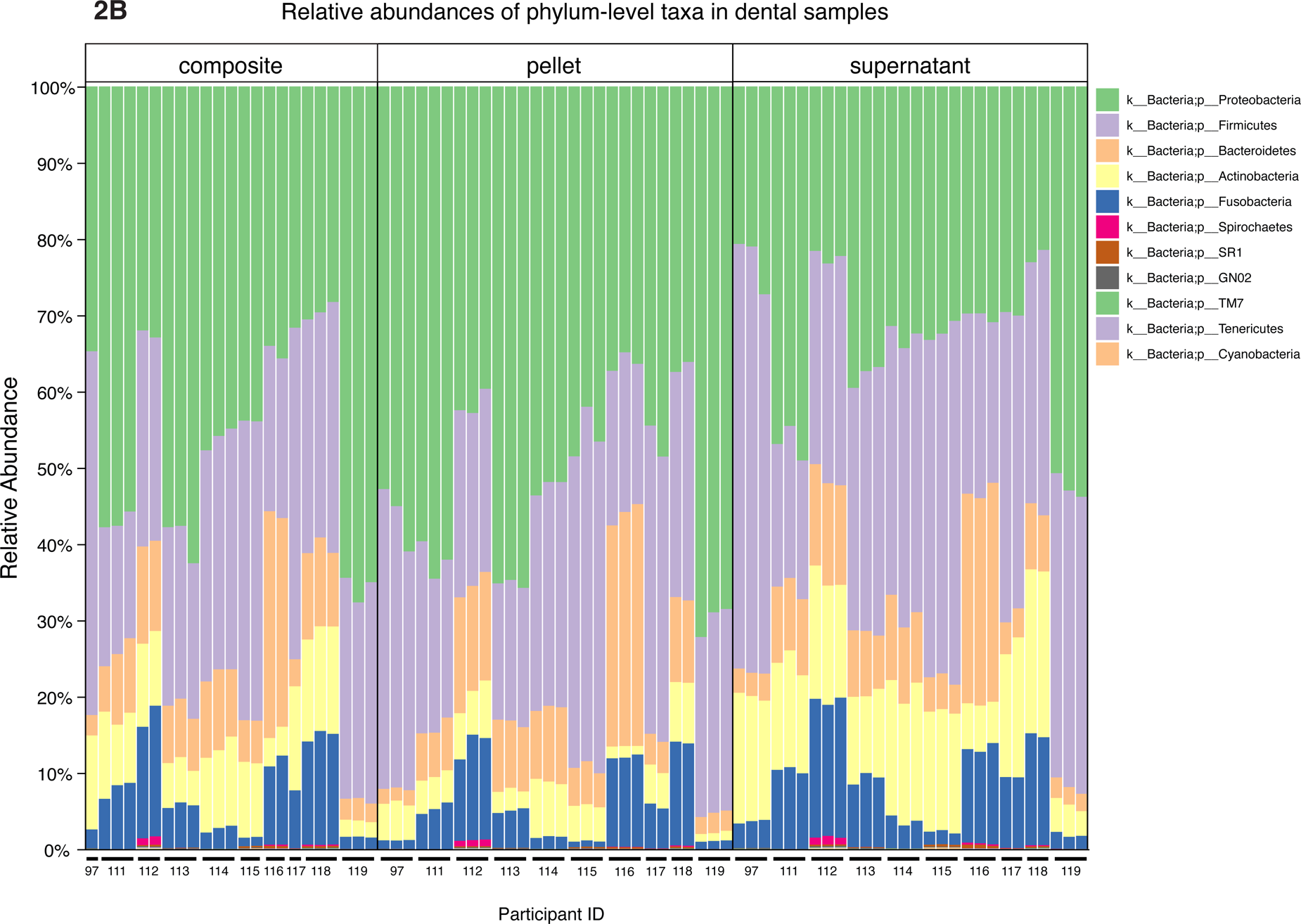
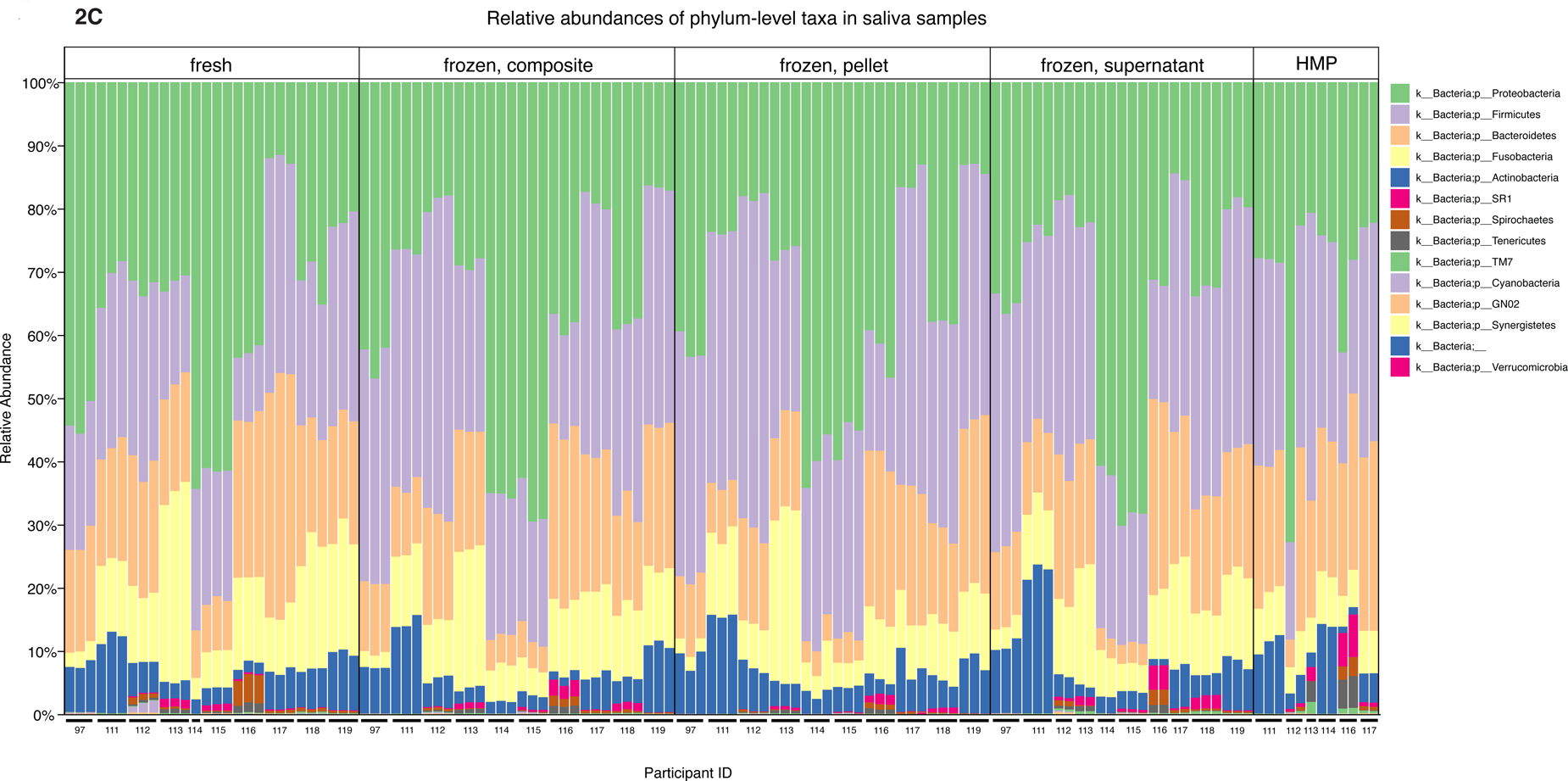
Microbial taxa differ at the phylum level for different fecal and oral sample processing methods. (A) Relative abundances of most prevalent microbial taxa of fecal samples categorized by treatment group (lyophilized, fresh, LN2, and LN2post72hr). (B) Relative abundances of most prevalent microbial taxa of dental samples categorized by thawed fraction (composite, pellet, and supernatant). (C) Relative abundances of most prevalent microbial taxa of saliva samples categorized by treatment (fresh, composite, pellet, supernatant, and HMP). (A-C) All legends list taxa in order of most to least abundant. (A-C) Taxa that were not visible due to low relative abundances are not listed in the figure key.

Relative abundances of phyla observed in oral samples are shown in Figure 2B and Figure 2C, with the top two dominant phyla being Proteobacteria and Firmicutes for both saliva and dental samples. Dental supernatant contained a higher relative abundance of Firmicutes compared to composite and pellet, but a lower relative abundance of Proteobacteria compared to composite and pellet (p<0.0001 for all). Dental pellet also contained a higher relative abundance of Proteobacteria compared to composite (p<0.0001). The relative abundance of Firmicutes, but not Proteobacteria, was significantly affected by treatment type in saliva samples. Fresh saliva contained a lower relative abundance of Firmicutes than composite (p<0.0001), HMP (p=0.002), pellet (p<0.0001), and supernatant (p<0.0001). Pellet contained a higher relative abundance of Firmicutes compared to composite (p=0.03) and HMP (p=0.01).

### Lyophilization strongly affects beta diversity of fecal samples

To compare microbial communities between samples, we calculated unweighted (taking into account presence/absence of microbes only) and weighted (also accounting for relative abundances of microbes present) UniFrac distances as measures of beta diversity. We then visualized these distances using Principal Coordinates Analysis (PCoA) and calculated whether groups of samples were different from each other by performing permutational multivariate analysis of variance (PERMANOVA). We found that donor was the primary driver of fecal sample clustering using unweighted UniFrac distances, explaining 77% of the variation in distances between samples (R^2^ = 0.77, F=39.98, p=0.001) (Figure 3A). However, treatment still explained a statistically significant amount of the variation (R^2^ =0.029, F=1.14, p=0.005) (Figure 3A). Samples were more strongly clustered by treatment based on weighted UniFrac compared to unweighted UniFrac distances (Figure 3B). Using weighted distances, lyophilized samples clustered away from the other treatments (R^2^=0.27, F=14.58, p=0.005), and donor identity explained less of the variation between distances than with unweighted distances (R^2^=0.59, F=17.48, p=0.001) (Figure 3B). For both unweighted and weighted UniFrac distances, we then performed pairwise PERMANOVA analyses, stratified by donor, to test which treatment types were significantly different from one another within each participant. We found that between-treatment distances were significantly greater than within-treatment distances for all treatment types (p<0.05), except for LN2 vs LN2post72hr samples. However, when using weighted UniFrac distances, lyophilization had a strong effect such that the lyophilized sample cluster was significantly different from all the other treatments even without stratification by donor (p=0.001 for all pairwise comparisons with lyophilized) (Figure 3B, Figure 3C).

**Figure 3.**
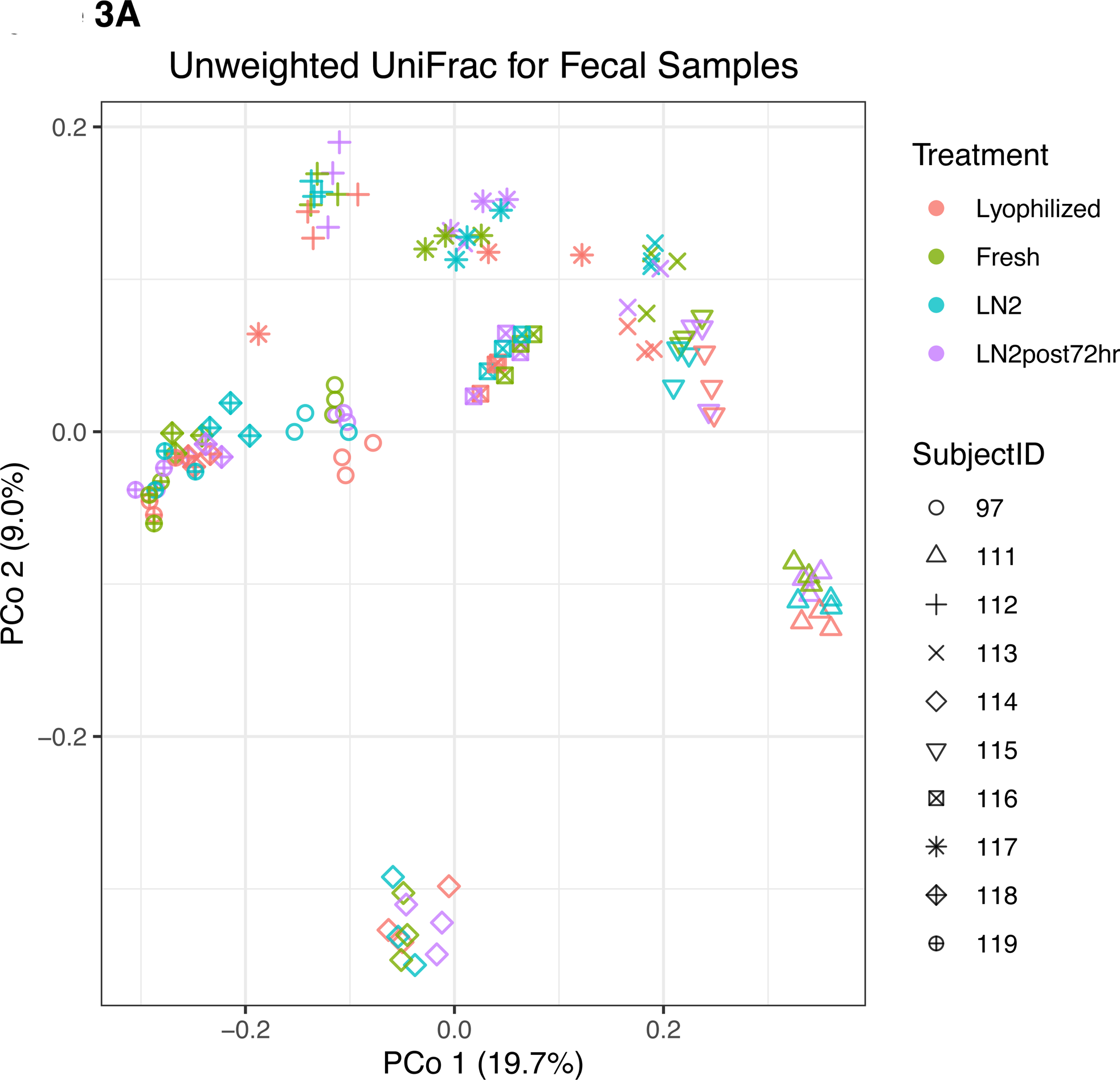
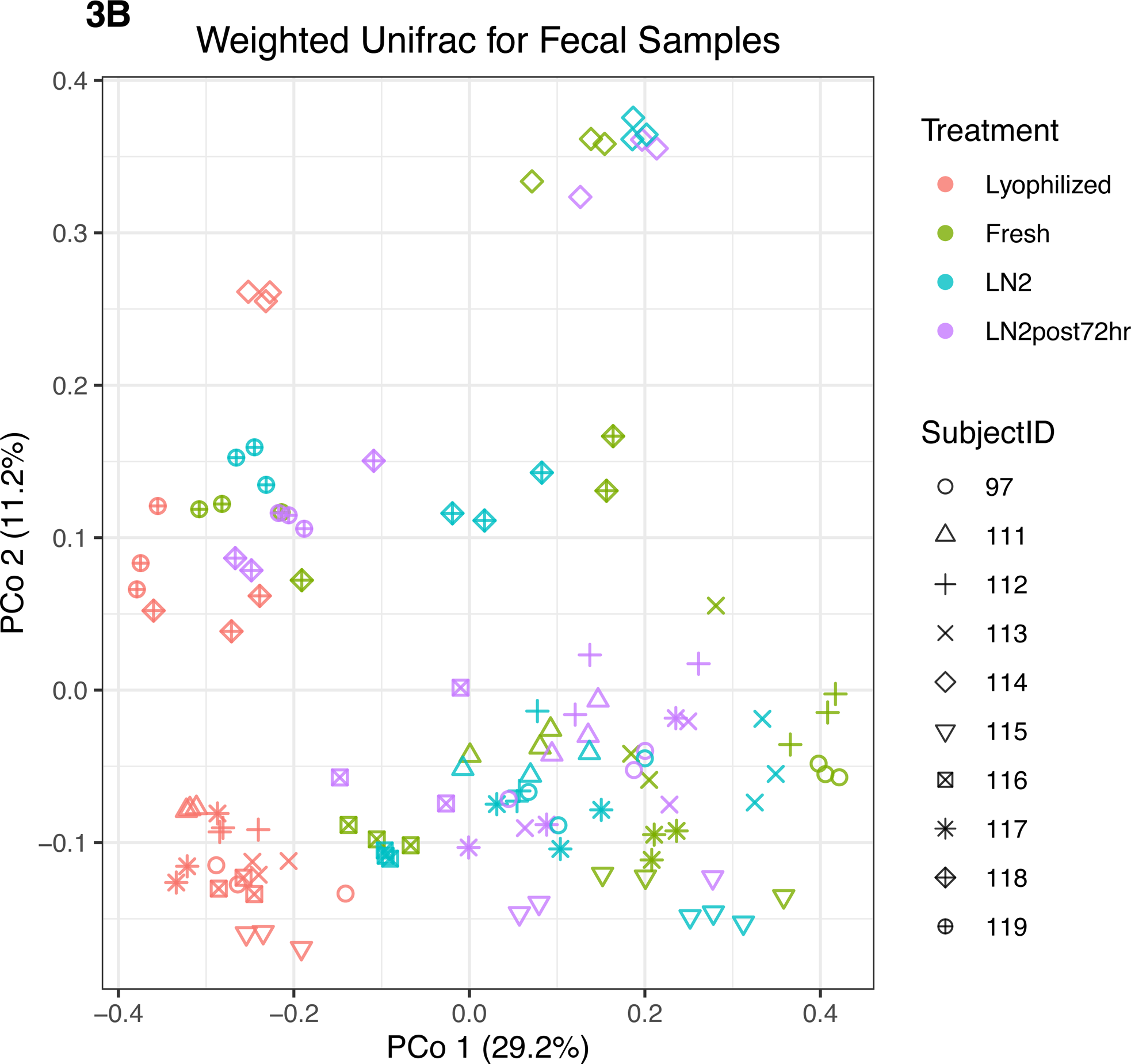
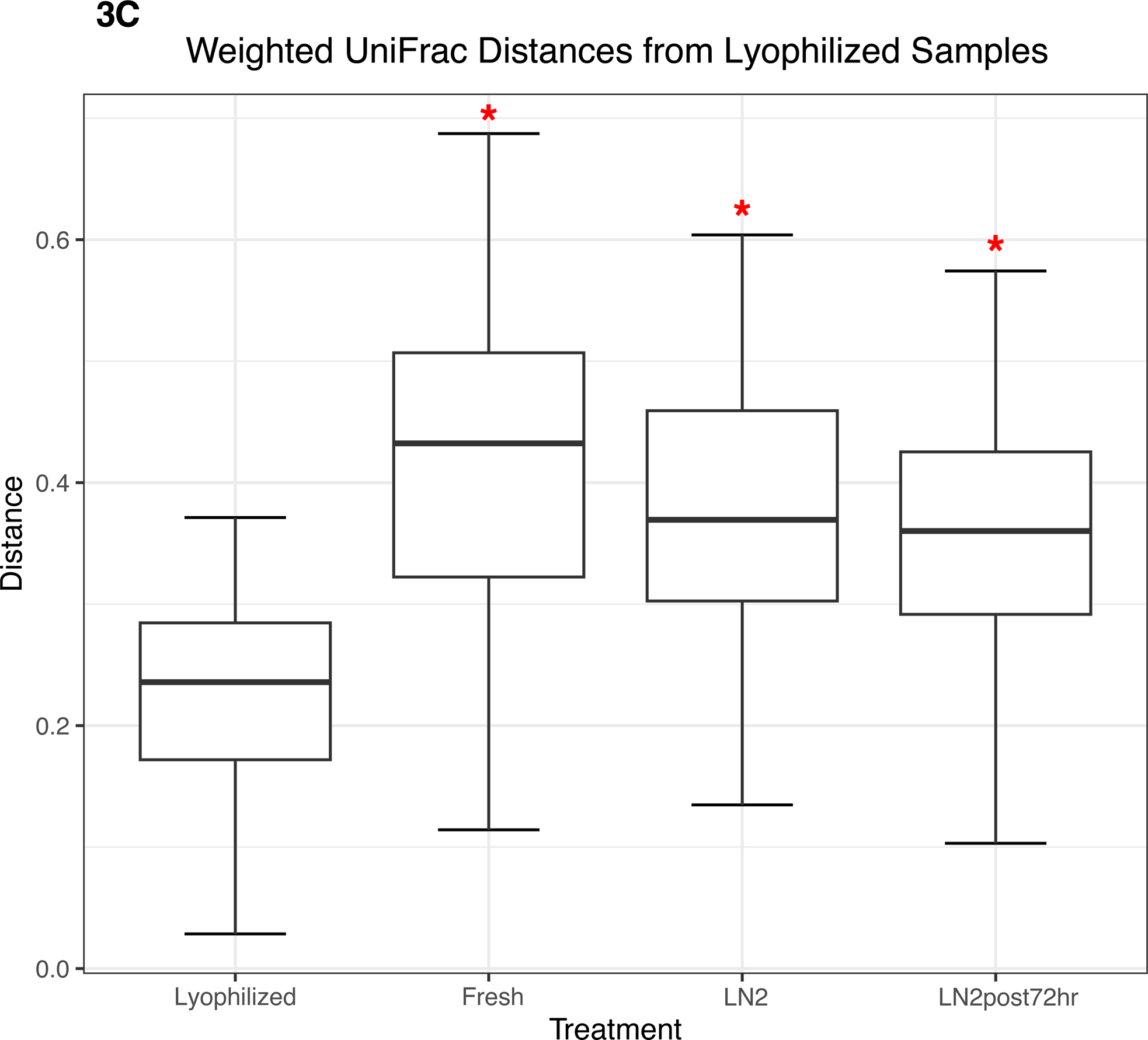
Beta diversity of fecal samples. PCoA plot of (A) unweighted and (B) weighted UniFrac distances for fecal samples, colored by treatment. Shapes represent different subjects. (C) Box plot representing pairwise weighted UniFrac distances of all treatment groups from lyophilized fecal samples. Red asterisks (*) indicate significantly different distances from lyophilized samples as determined by PERMANOVA without stratification by participant.

### Treatment method affects beta diversity of dental samples

Next, we compared beta diversity for dental samples obtained from thawed fractions. Using unweighted UniFrac distances, we found donor identity explained most of the variation between samples (R^2^=0.73, F=21.19, p=0.001), with visually apparent clustering (Figure 4A). Overall, fraction had a small yet statistically significant effect (R^2^=0.02, F=0.70, p=0.005). Using weighted UniFrac distances, donor identity explained more of the variation between samples (R^2^=0.83, F=37.46, p=0.001) as compared to when using unweighted UniFrac distances, while fraction also explained more of the variation observed (R^2^=0.14, F=5.93, p=0.005) compared to unweighted distances (Figure 4B). We then performed a PERMANOVA stratified by donor using weighted UniFrac distances and found that all dental fractions were significantly different from one another (p=0.001 for all pairwise comparisons). However, when using weighted UniFrac distances and considering all samples collectively without stratifying by donor, we found that only supernatant samples were significantly different from pellet (p=0.001) and composite (p=0.009) (Figure 4C).

**Figure 4.**
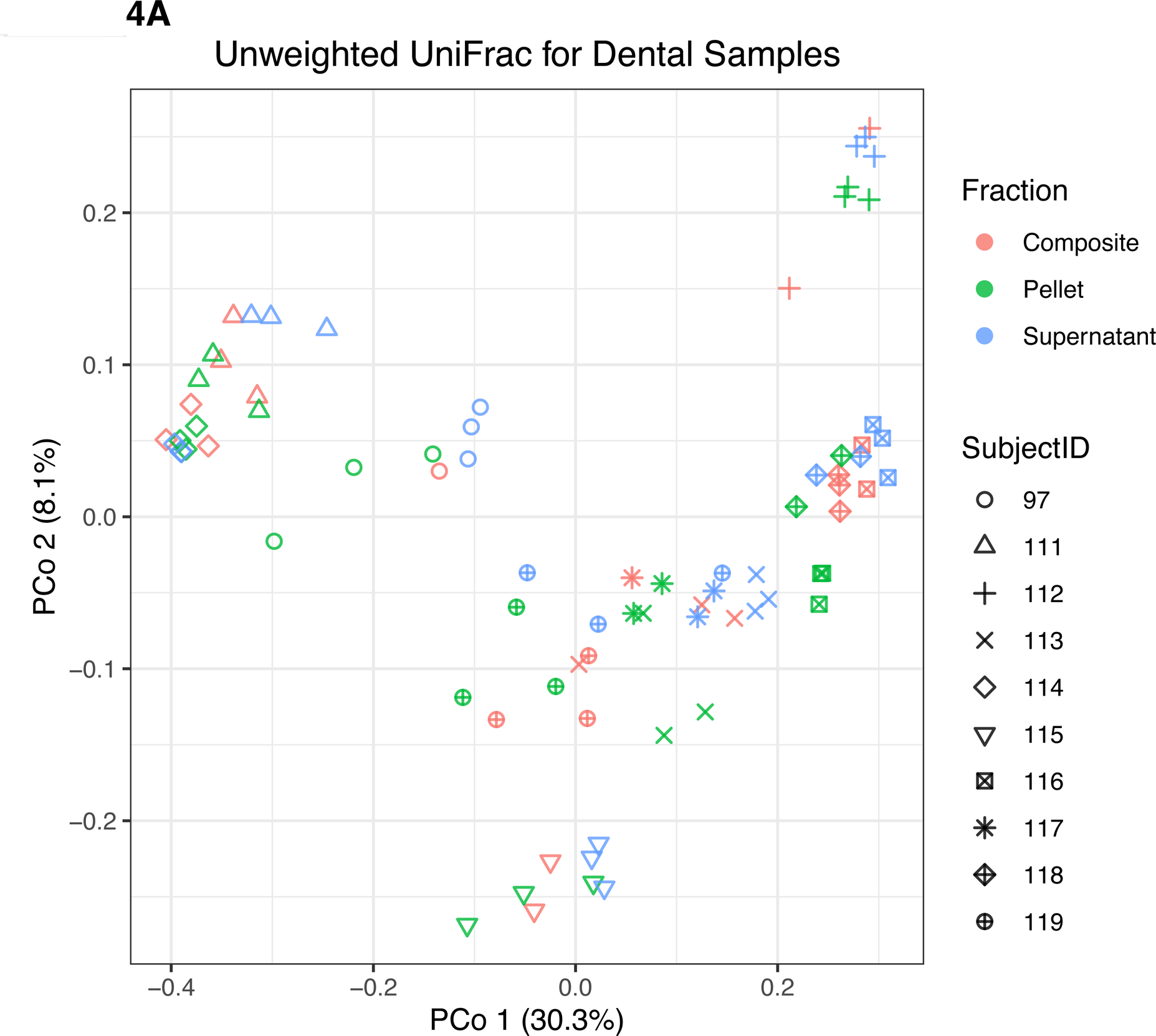
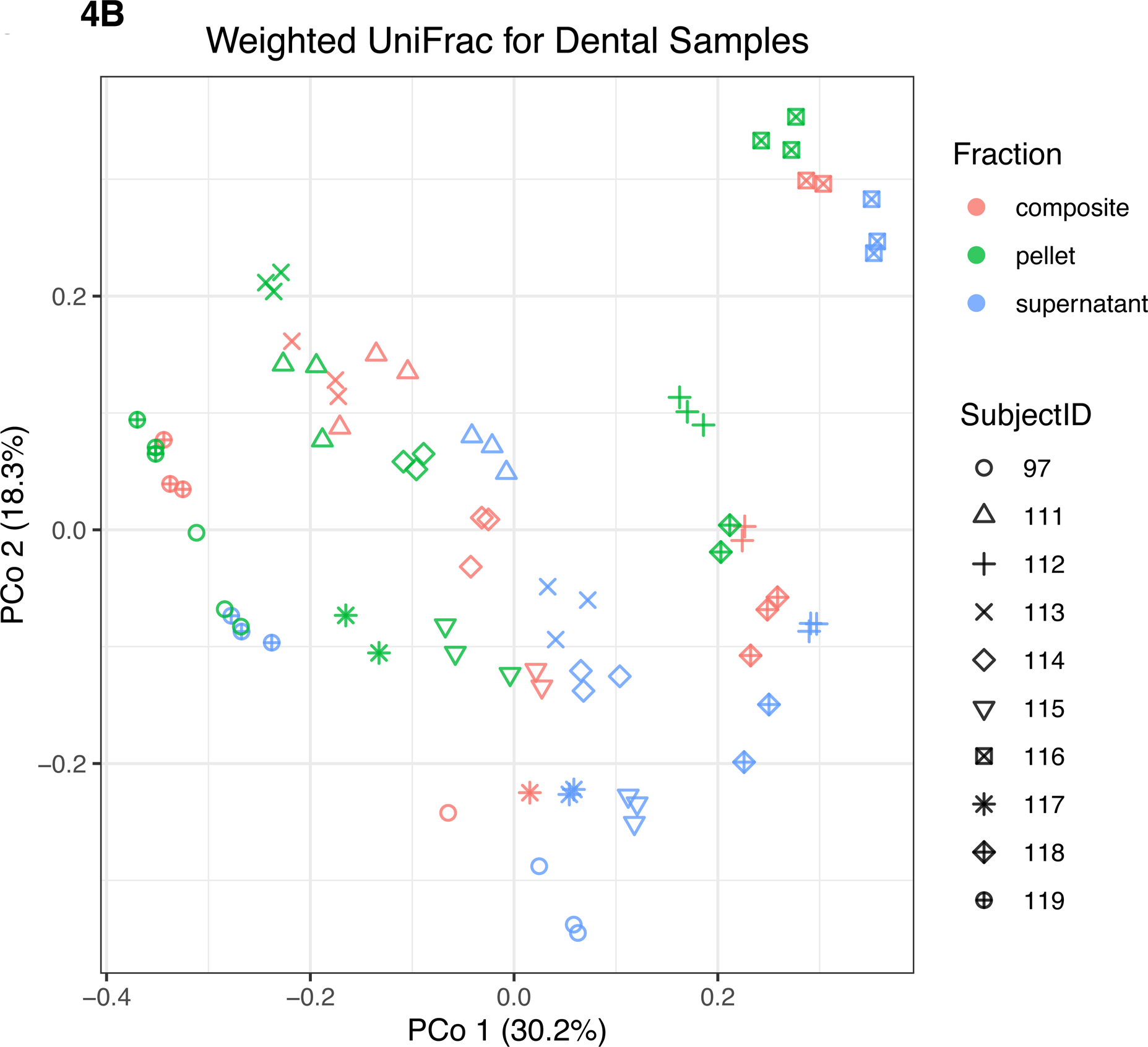
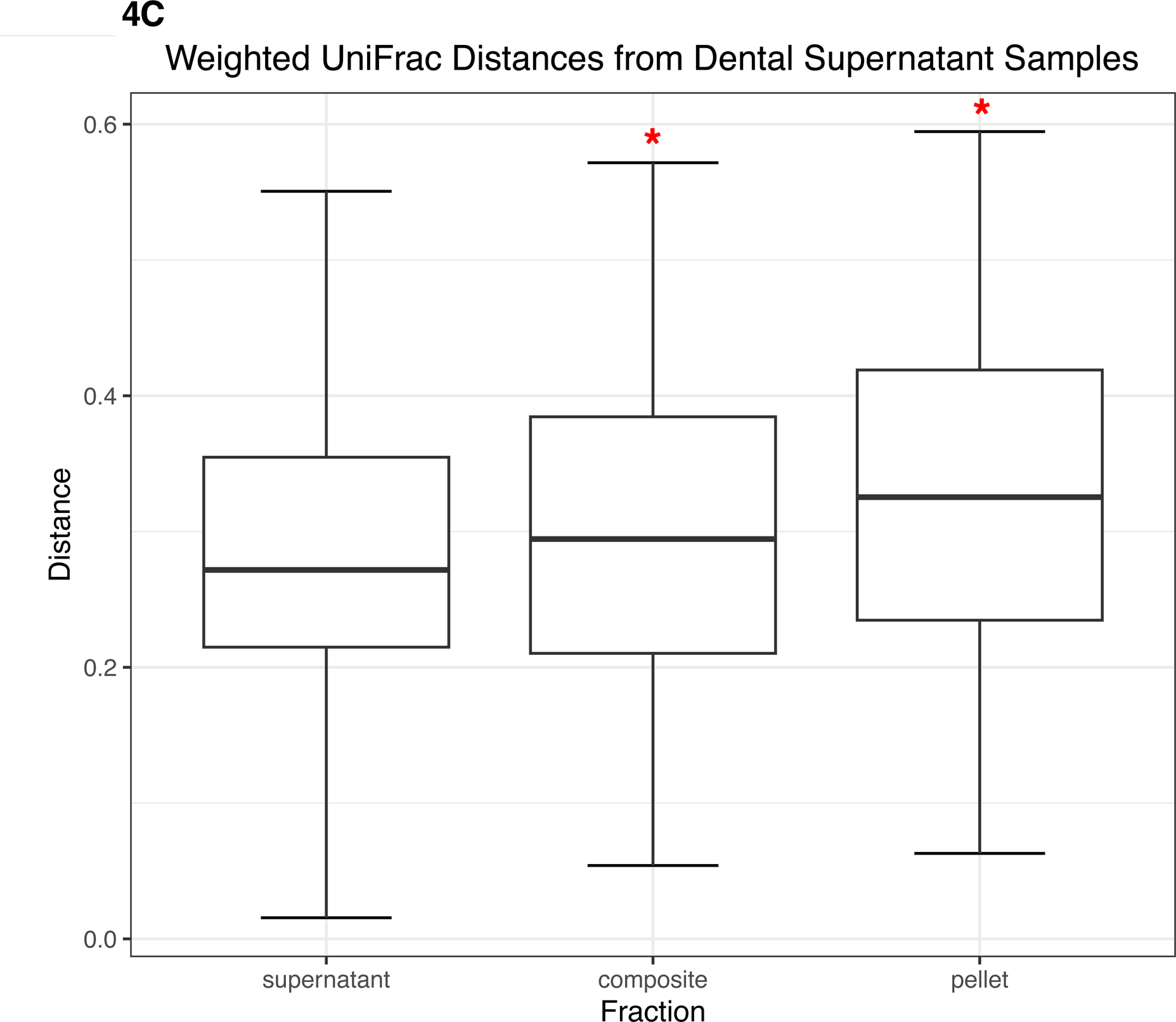
Beta diversity of dental samples. Principal coordinates analysis plot of (A) unweighted and (B) weighted UniFrac distances for dental samples, colored by fraction. Shapes represent different subjects. (C) Box plots represent pairwise weighted UniFrac distances of all dental fractions from dental supernatant. Red asterisks (*) indicate significantly different distances from dental supernatant samples as determined by PERMANOVA without stratification by participant.

### Treatment method affects beta diversity of saliva samples

We also compared beta diversity between different fractions of thawed saliva to fresh saliva samples. Samples from the HMP processing method were excluded from this analysis due to fewer replicates with adequate sequencing depth, which would have resulted in unbalanced groups in the statistical models. Using unweighted UniFrac distances, similar to our dental sample results, we found that donor identity explained most of the variation between saliva samples (R^2^=0.73, F=30.47, p=0.001), and treatment (which included composite, pellet, supernatant, and fresh) was a minor but still statistically significant contributor (R^2^=0.078, F=3.09, p=0.005) (Figure 5A). We also found that using weighted UniFrac distances increased the contribution of donor identity (R^2^=0.83, F=54.16, p=0.001) and decreased the contribution of treatment (R^2^=0.068, F=2.64, p=0.005) (Figure 5B). Despite the minor contribution of treatment relative to donor in explaining microbial community variation in saliva samples, when stratifying by donor, using both weighted and unweighted UniFrac distances, we found that all treatments had significantly greater between-sample type distances than within-sample type distances for all pairwise comparisons of treatment types (p=0.001 for all except composite vs supernatant using unweighted UniFrac which was p=0.048). Thus, although donor identity primarily separated microbial communities, treatment significantly affected microbial community differences between samples from the same donor. Without stratifying by donor, using both weighted and unweighted UniFrac distances, we found that only fresh samples were significantly different from the other treatment groups (p<0.05) (weighted UniFrac shown in Figure 5C).

**Figure 5.**
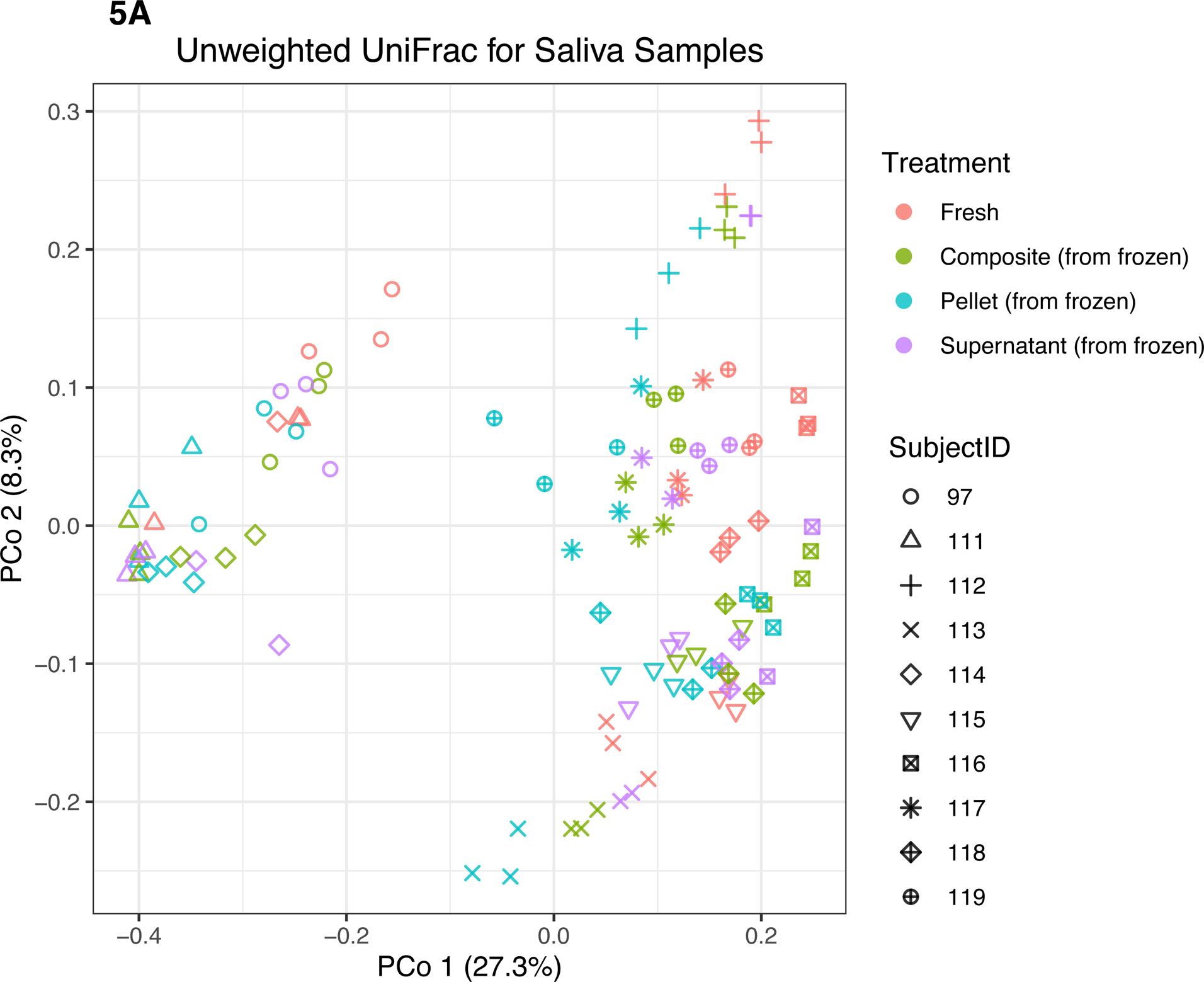
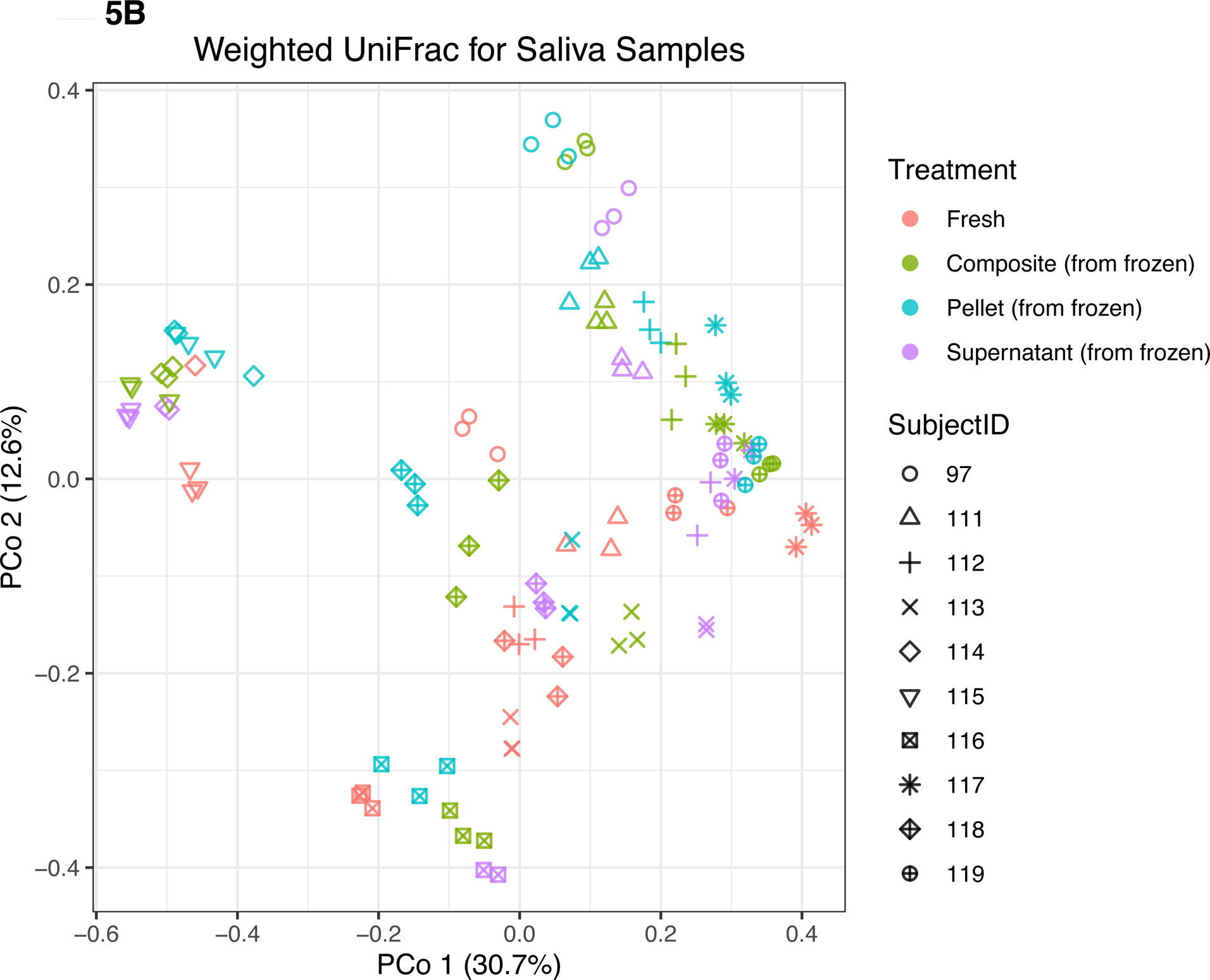
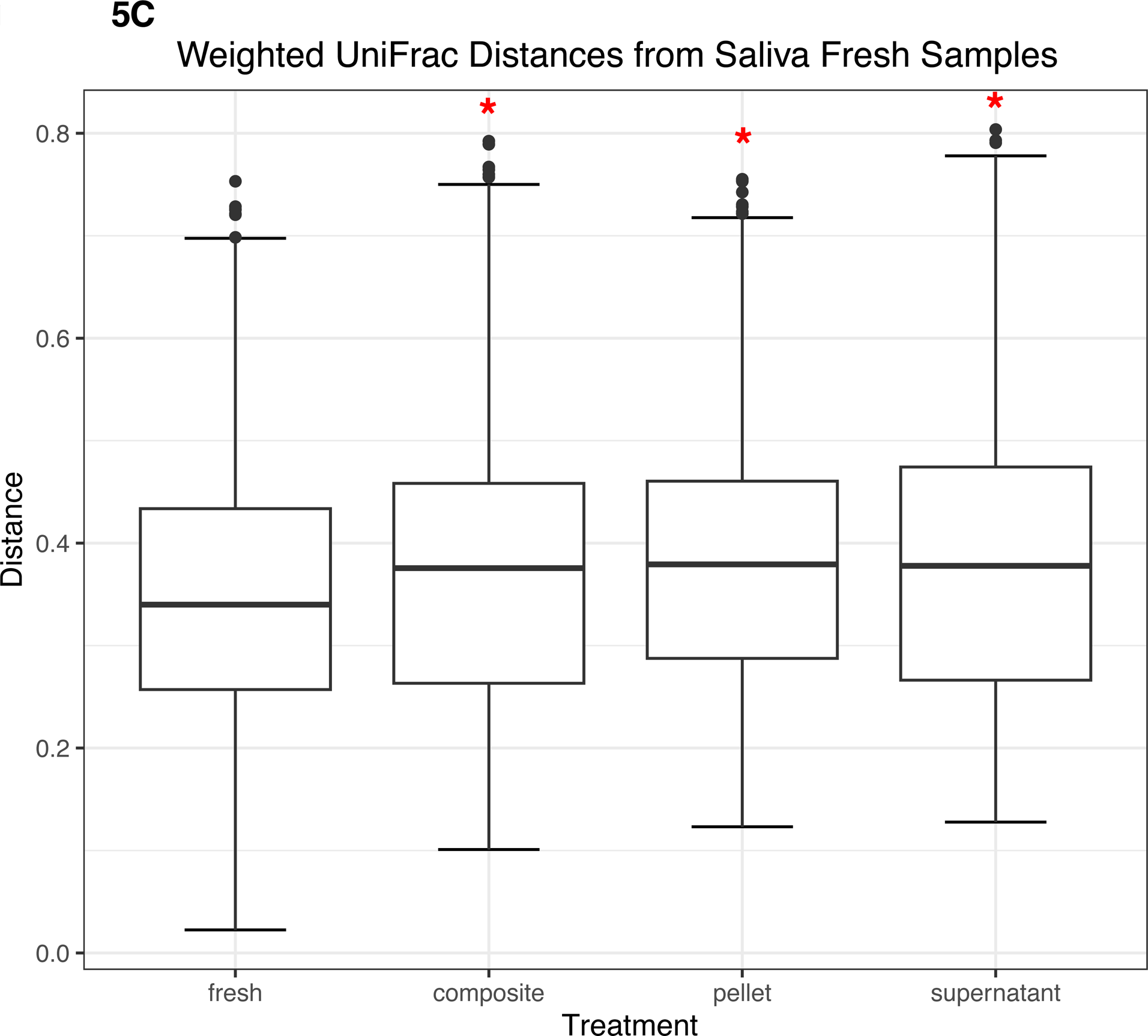
Beta diversity of saliva samples. Principal coordinates analysis plot of (A) unweighted and (B) weighted UniFrac distances for saliva samples, colored by treatment. Shapes represent different subjects. (C) Box plots represent pairwise weighted UniFrac distances of samples in each treatment group from fresh saliva samples. Red asterisks (*) indicate significantly different distances from fresh saliva samples as determined by PERMANOVA without stratification by participant.

### Alpha diversity is highest in lyophilized fecal samples, dental supernatant, and fresh saliva samples

In fecal samples, lyophilized samples had the highest alpha diversity overall. Despite no significant differences in the number of observed ASVs based on treatment type (Figure 6A), we found that lyophilized samples had significantly greater evenness than fresh (p<0.0001), LN2 (p=0.0005), and LN2post72hr samples (p=0.0007) (Figure 6B), and significantly greater Faith’s PD than fresh (p=0.03) and LN2 samples (p=0.01) (Figure 6C). Faith’s PD by individual donors is shown in Figure 6D. Interestingly, the LN2post72hr samples did not exhibit a greater range of standard deviation in Faith’s PD between replicates for each donor (SD=4.0) than the lyophilized samples (SD=5.8) even though the LN2post72hr samples were stored at room temperature for three days with no added preservative.

**Figure 6.**
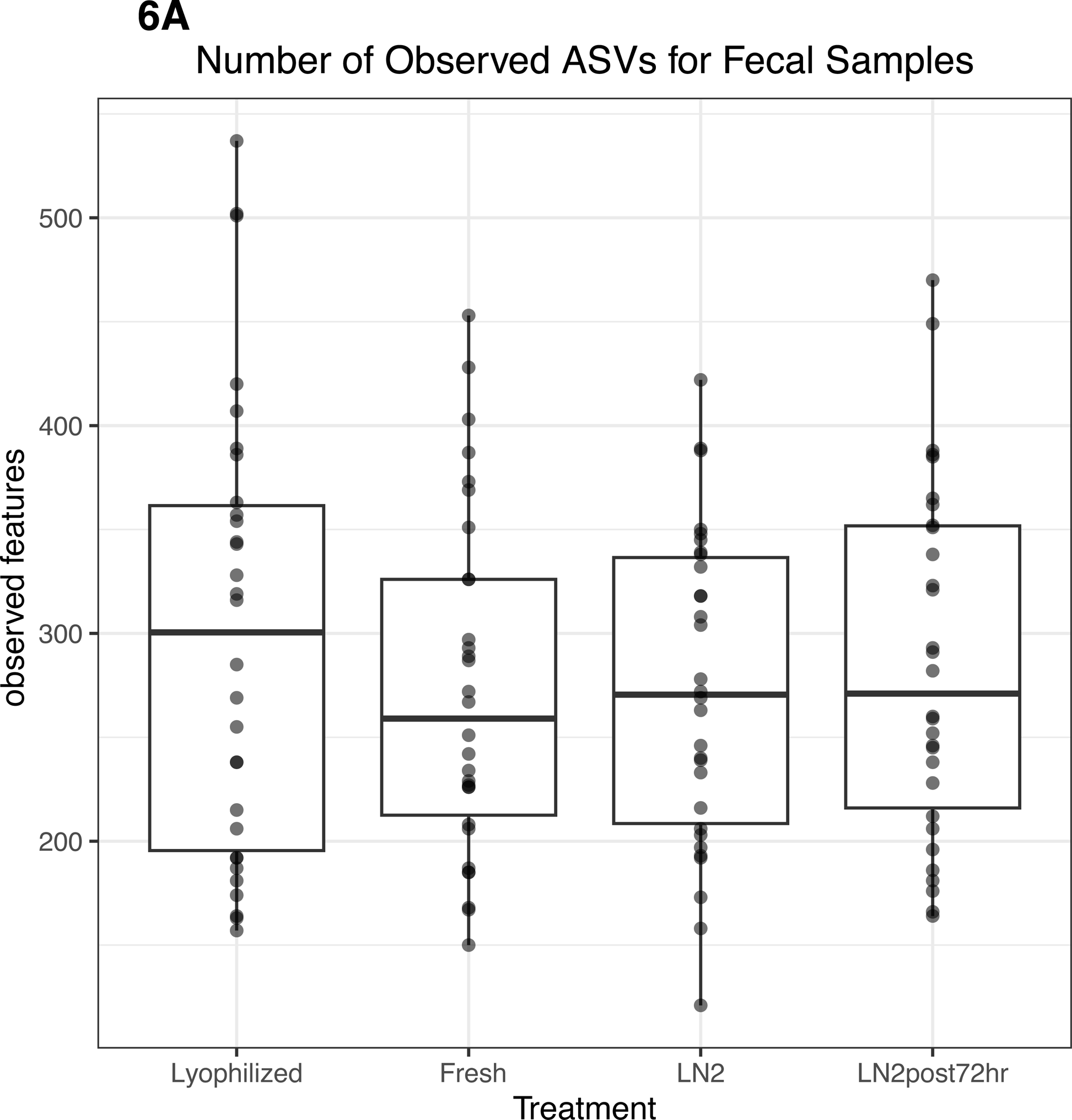
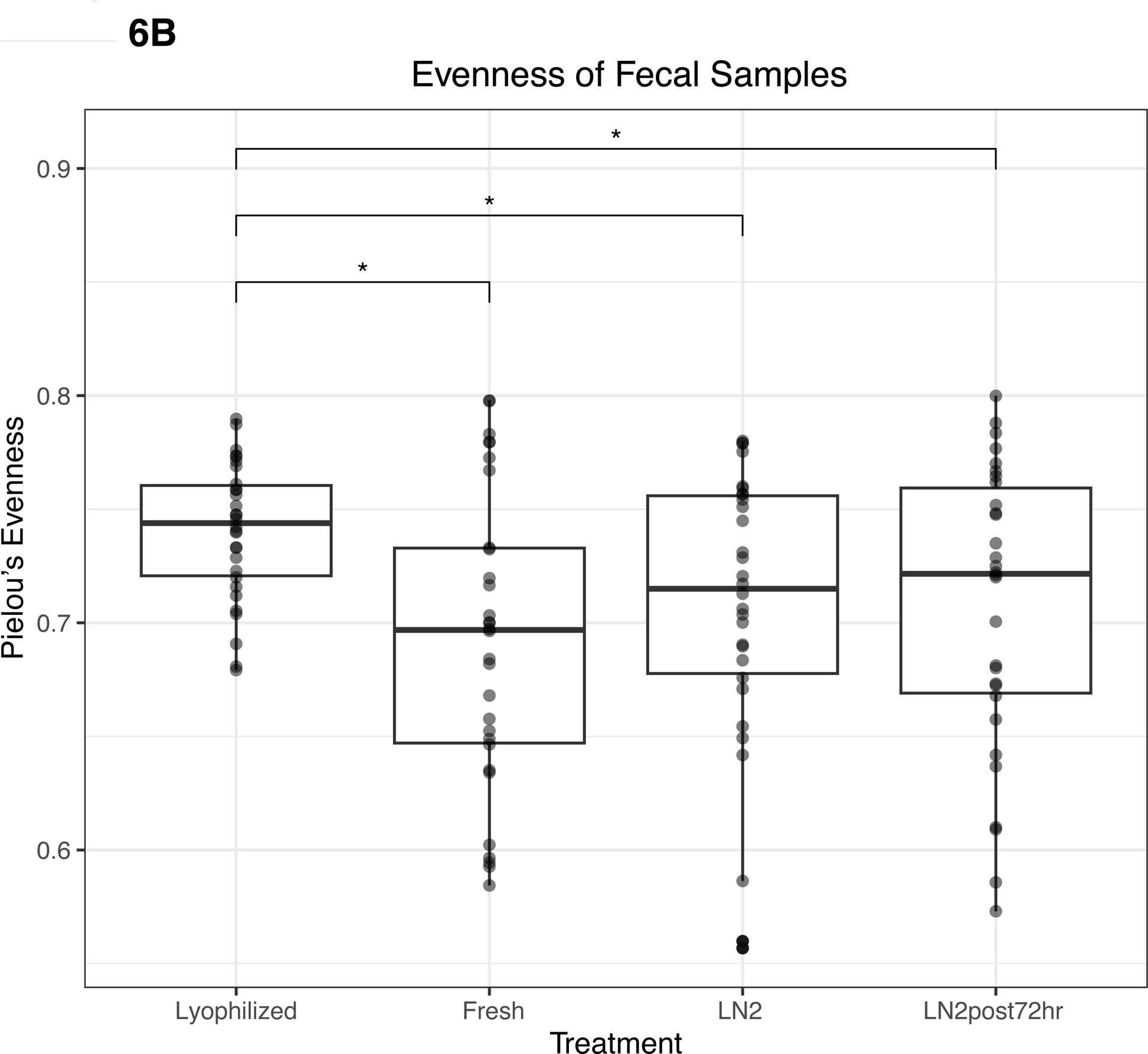
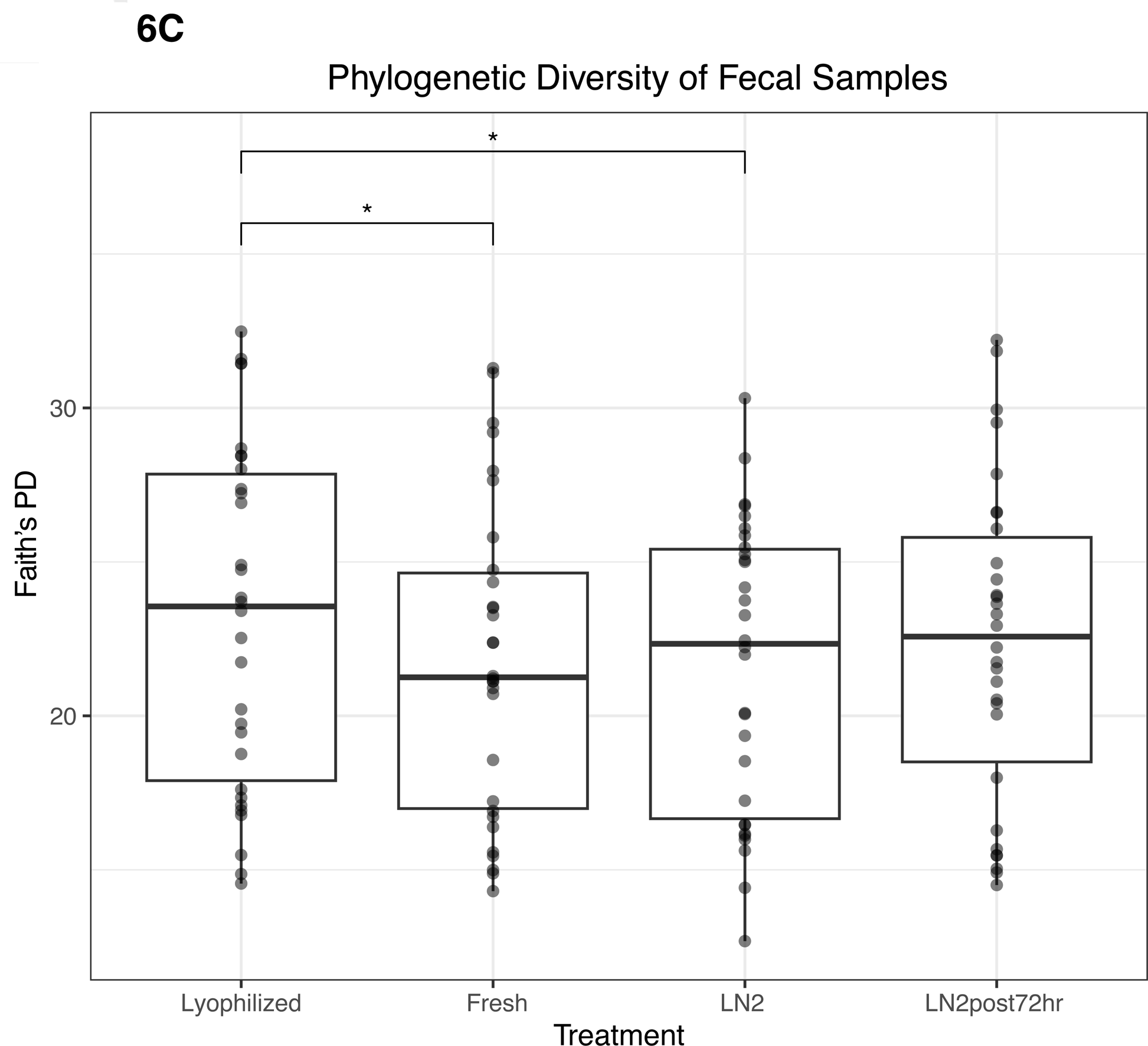
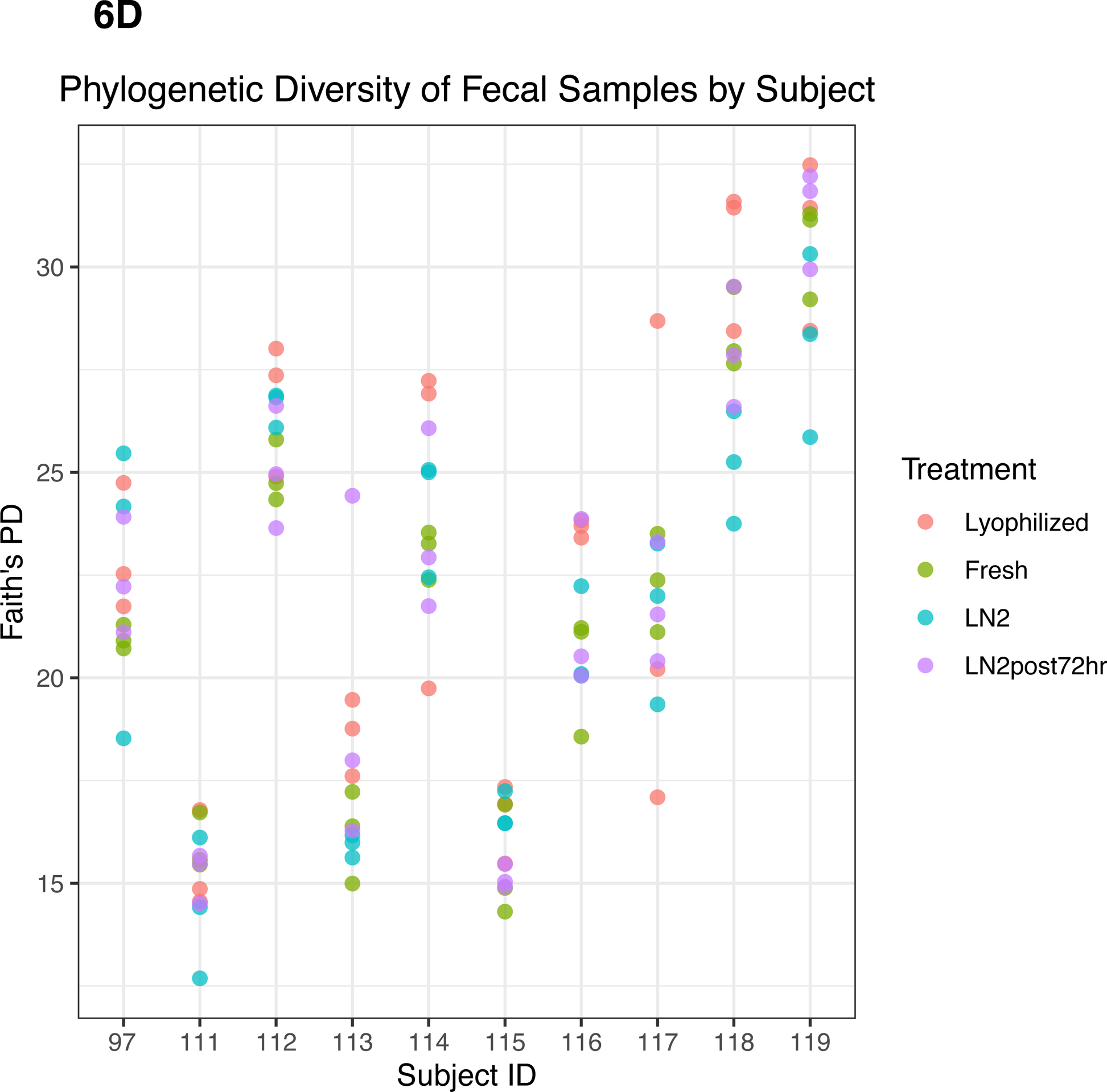
Alpha diversity of fecal samples. (A) Observed ASVs in fecal samples by treatment type. (B) Evenness of fecal samples by treatment type. (C) Phylogenetic diversity as determined by Faith’s PD of fecal samples by treatment type. Statistically significant (p<0.05) differences between groups as determined by pairwise comparisons are denoted by an asterisk (*) in A-C. (D) Phylogenetic diversity in individual subjects, colored by treatment type.

For dental samples, we found that the supernatant fraction had the highest alpha diversity. Supernatant had a greater number of observed ASVs than both composite (p=0.0004) and pellet (p<0.0001) fractions (Figure 7A). All fractions differed in evenness: supernatant had greater evenness than composite (p=0.0001) and pellet (p<0.0001), and composite had greater evenness than pellet (p<0.0001) (Figure 7B). Finally, supernatant had greater Faith’s PD than both composite (p=0.02) and pellet (p=0.0001) (Figure 7C). Faith’s PD in donor replicates is shown in Figure 7D.

**Figure 7.**
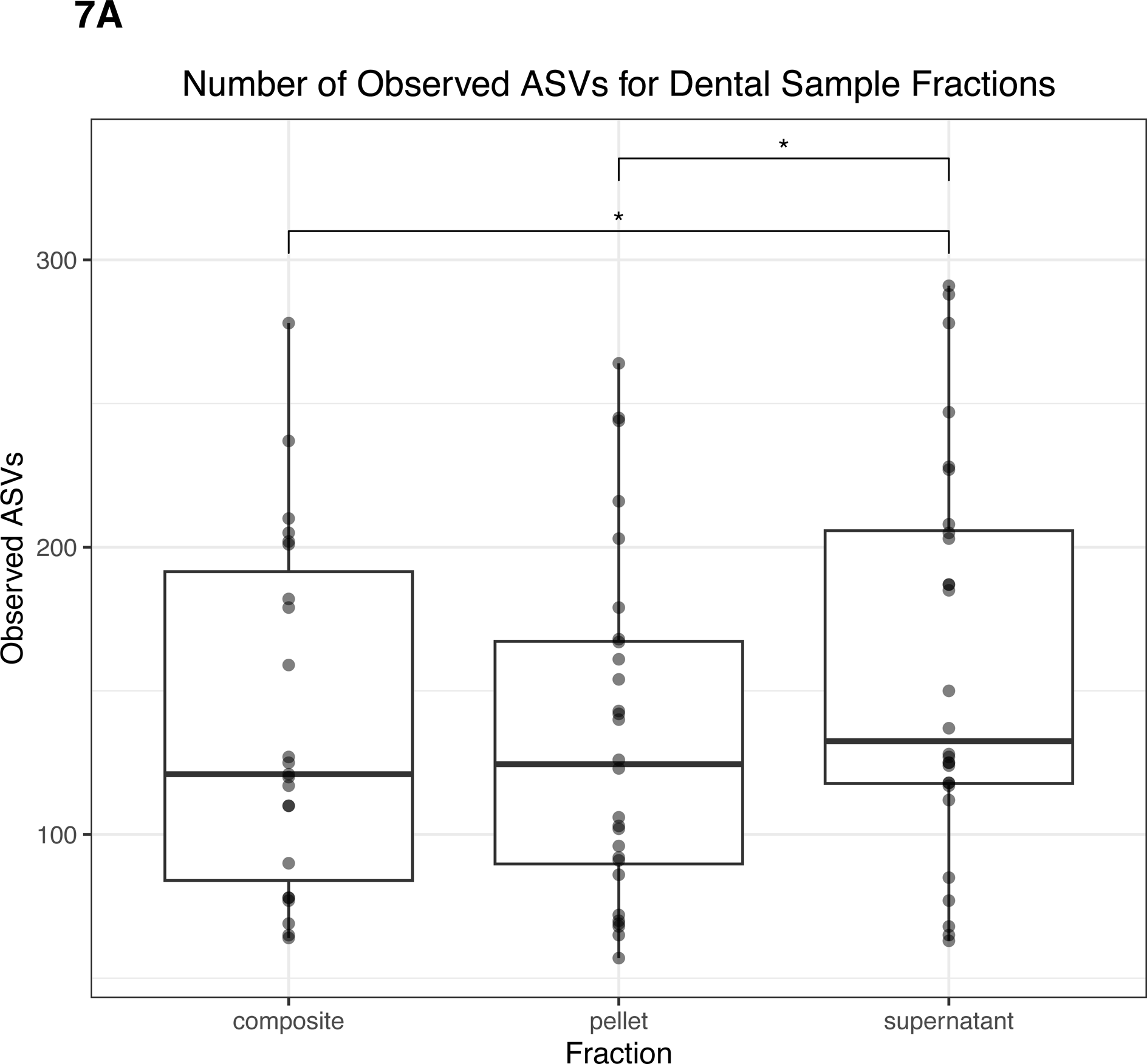
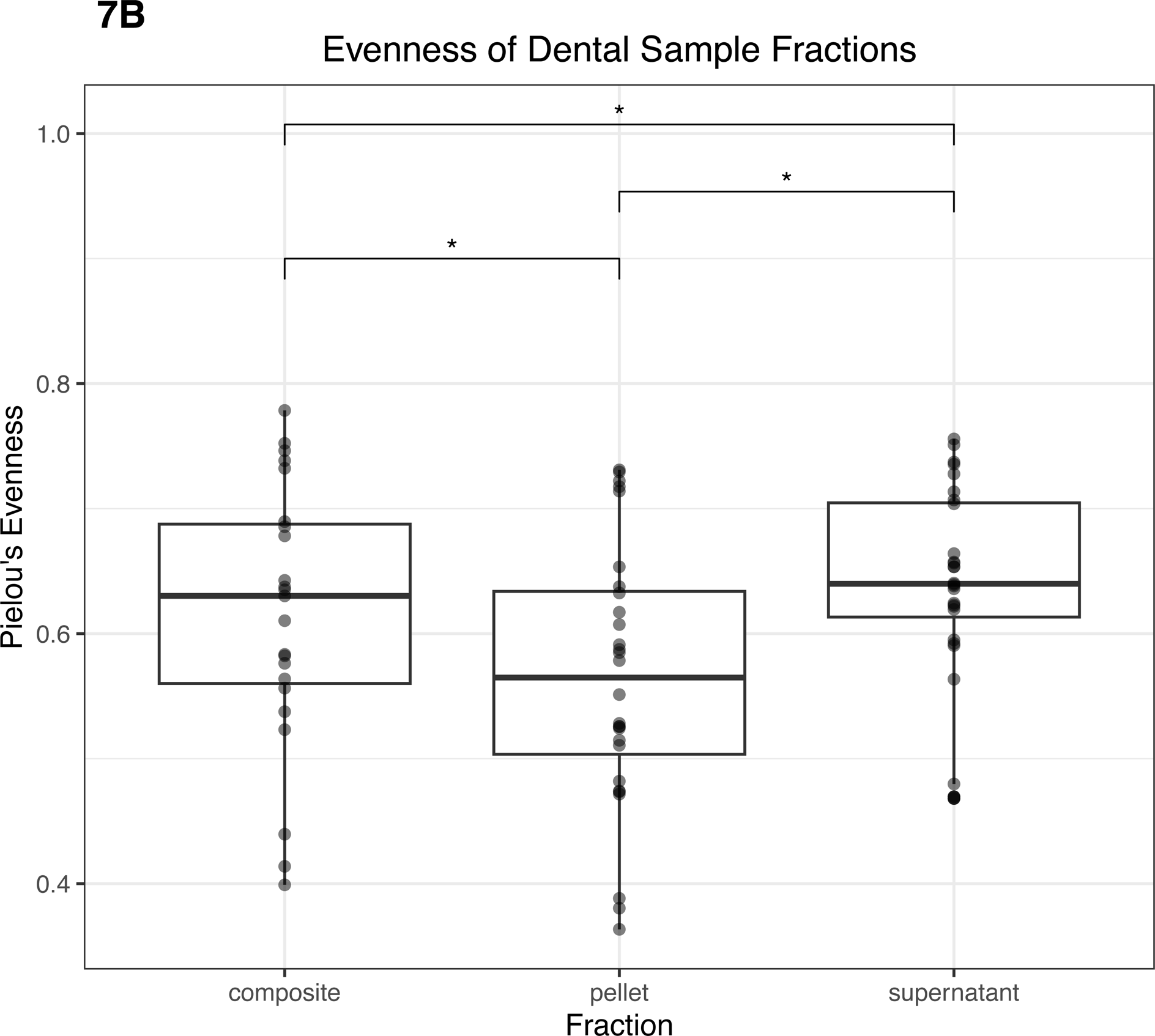
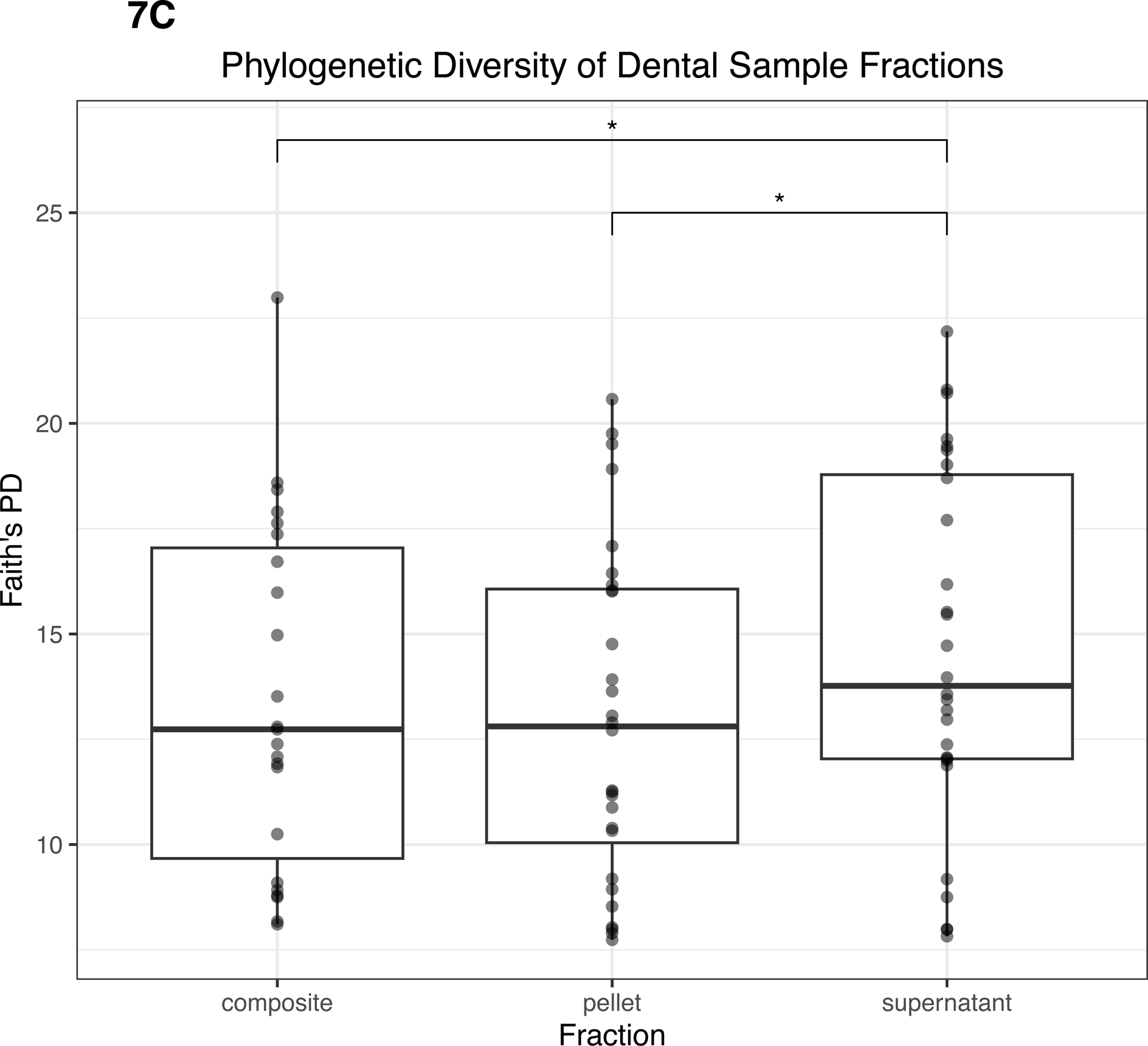
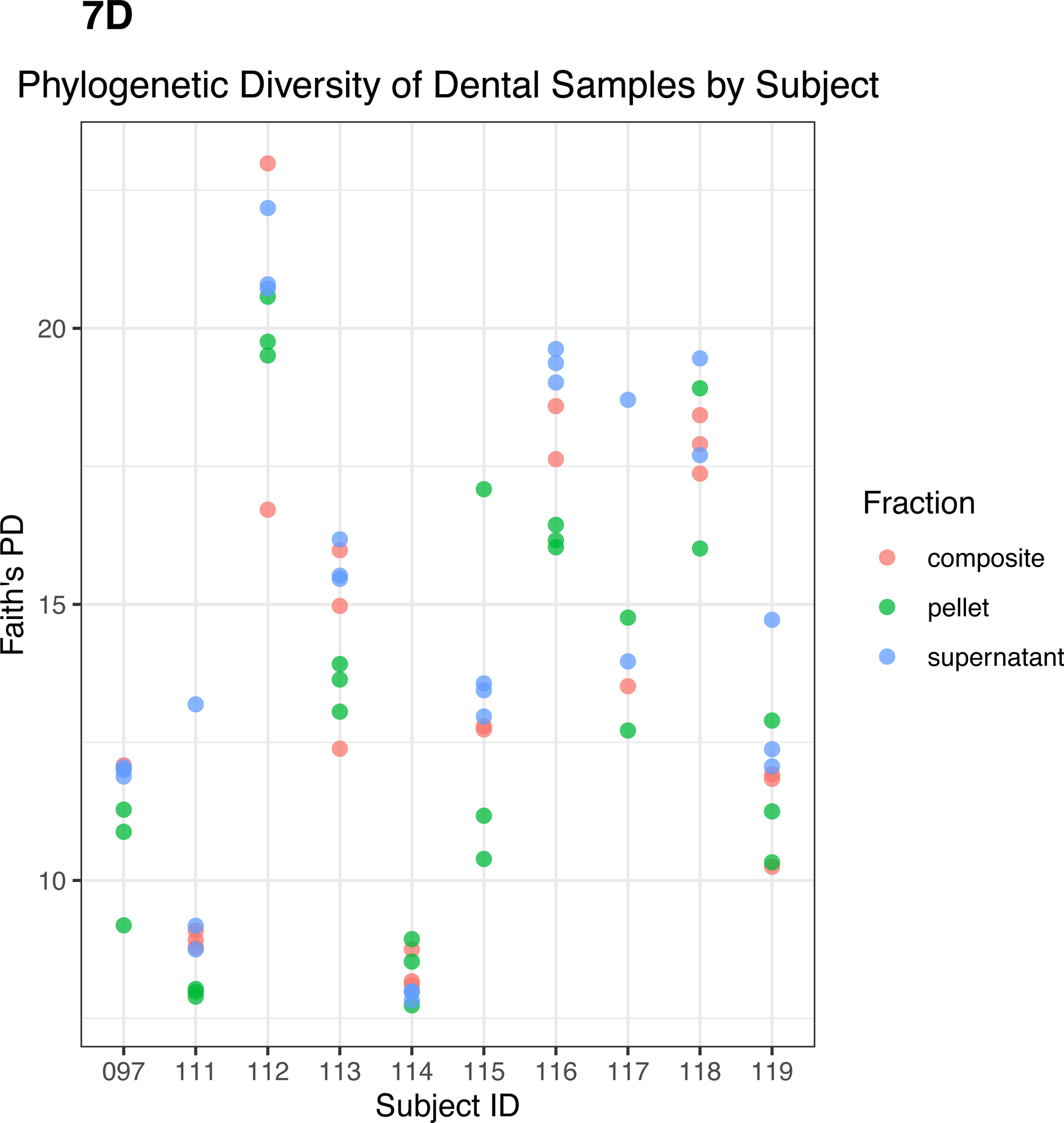
Alpha diversity of dental samples. (A) Observed ASVs in dental samples by fraction. (B) Evenness of dental samples by fraction. (C) Phylogenetic diversity as determined by Faith’s PD of dental samples by fraction. Statistically significant (p<0.05) differences between fractions are denoted by an asterisk (*) in A-C. (D) Phylogenetic diversity by individual subject, colored by dental fraction type.

For saliva samples, we found that fresh saliva had the highest alpha diversity. Fresh saliva samples had a higher number of observed ASVs than all thawed fractions (p<0.05) (Figure 8A). Composite and supernatant both had a greater number of observed ASVs than pellet (p<0.05) (Figure 8A). Fresh saliva had greater evenness than pellet (p=0.003) and composite (p=0.02), and supernatant had greater evenness than pellet (p=0.0003) and composite as well (p=0.002) (Figure 8B). However, there were no significant differences in evenness between fresh and supernatant or composite and pellet (p>0.05) (Figure 8B). Fresh saliva samples had the greatest Faith’s PD compared to all frozen fractions (p<0.0001) (Figure 8C). While supernatant and composite were not significantly different from each other, they both had significantly greater Faith’s PD than pellet (p=0.01). Faith’s PD in donor replicates for each saliva sample type can be found in Figure 8D. Additionally, we found that the average Faith’s PD across saliva samples and dental samples was correlated in donors (r=0.88, p=0.0009).

**Figure 8.**
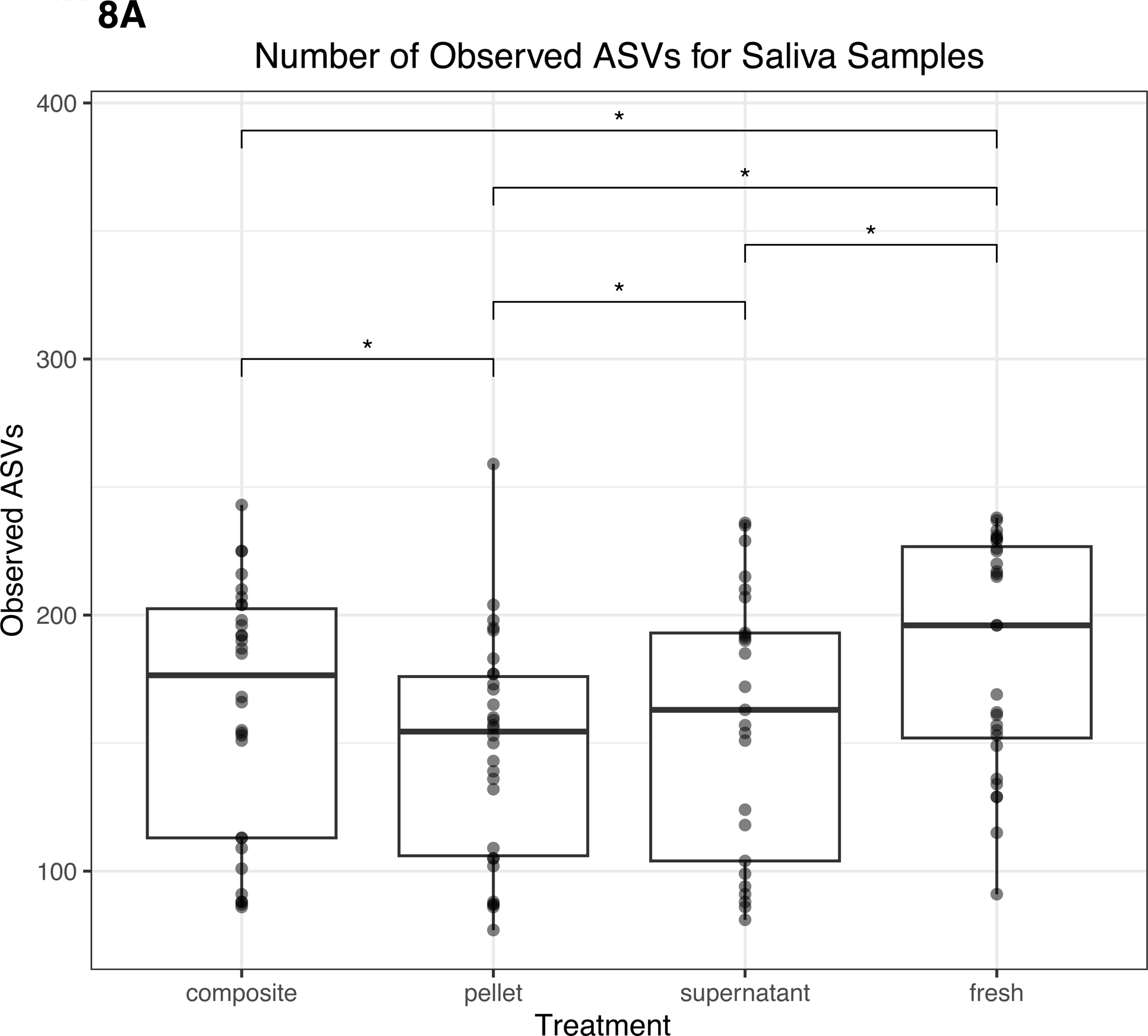
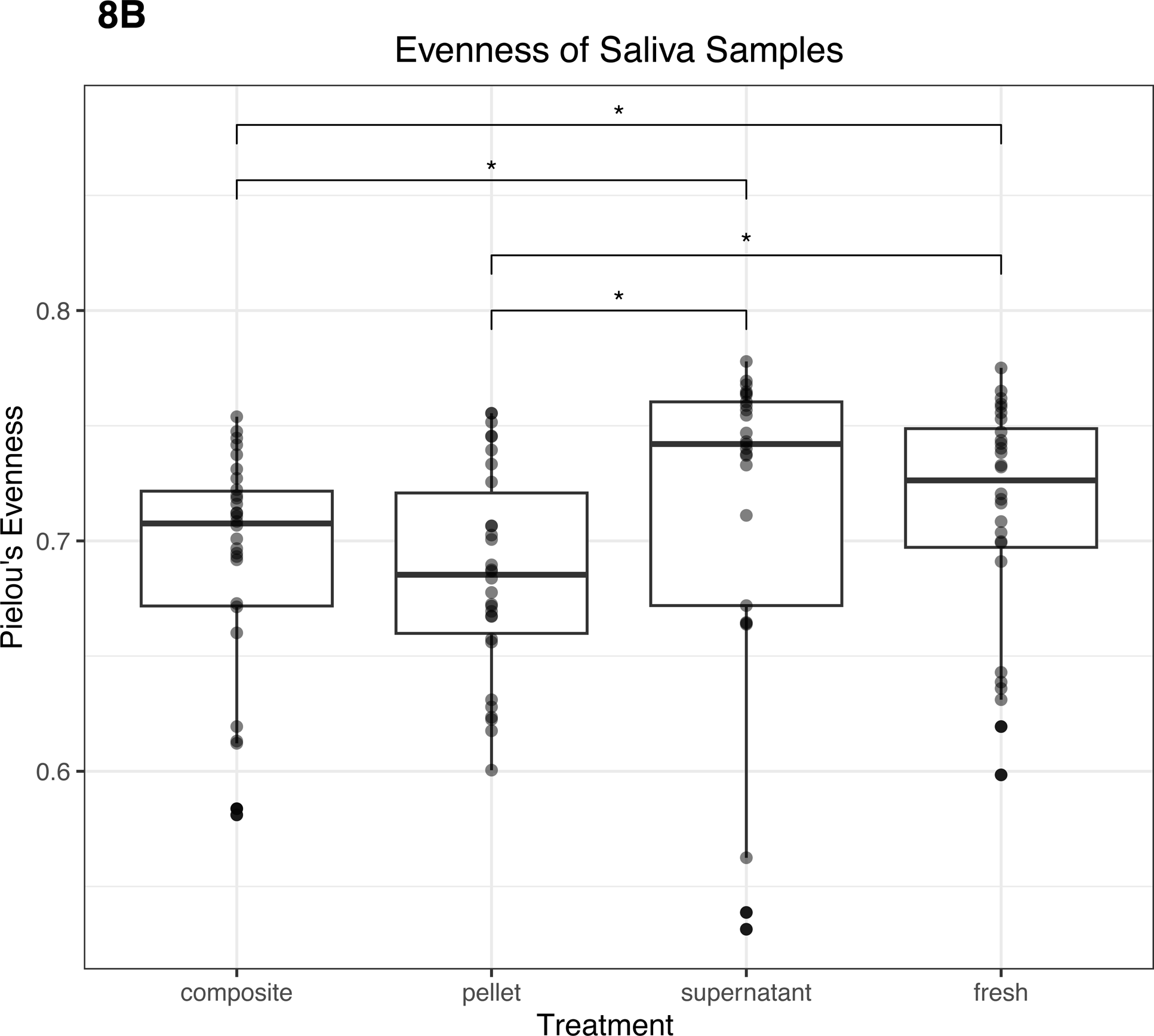
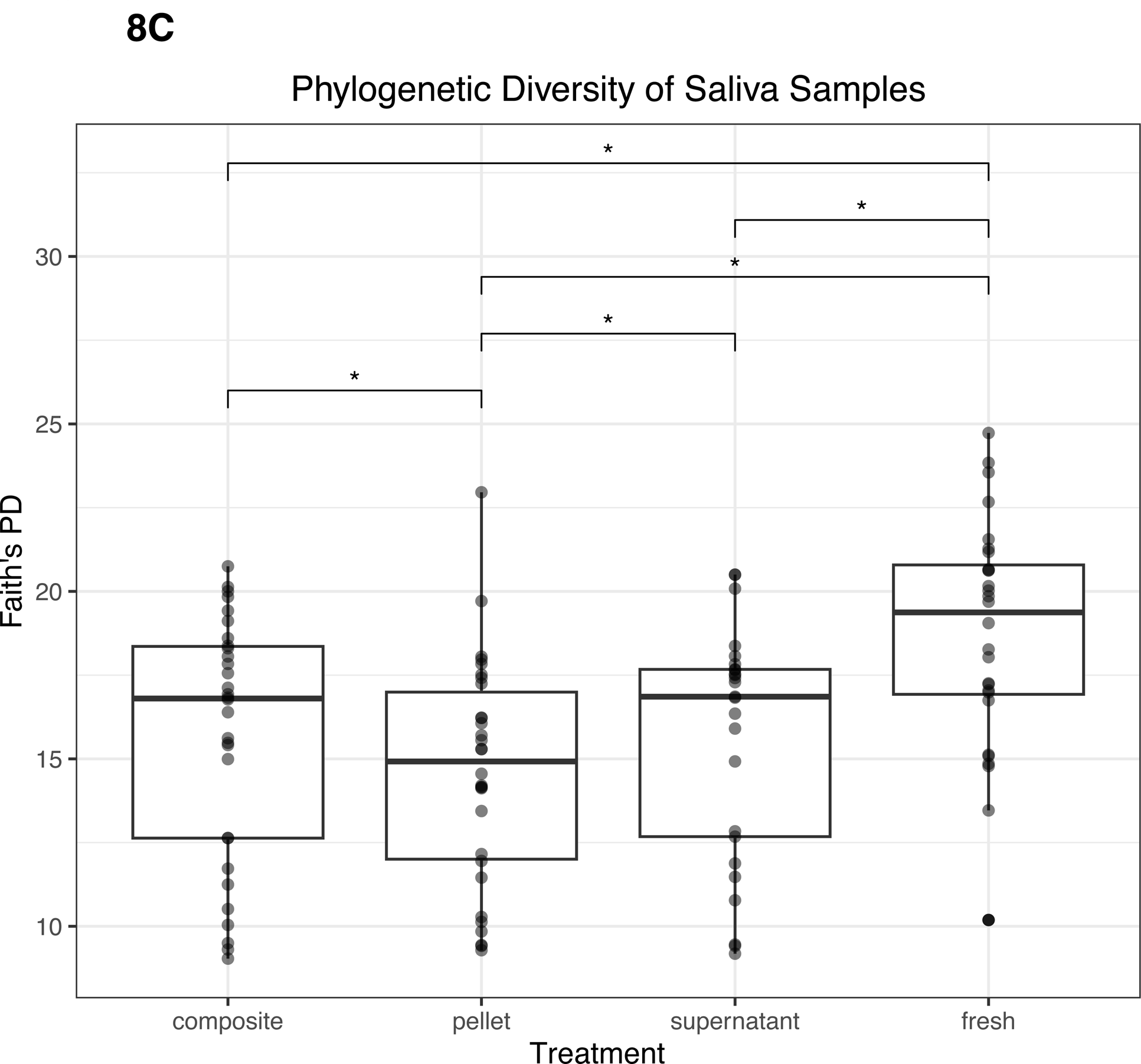
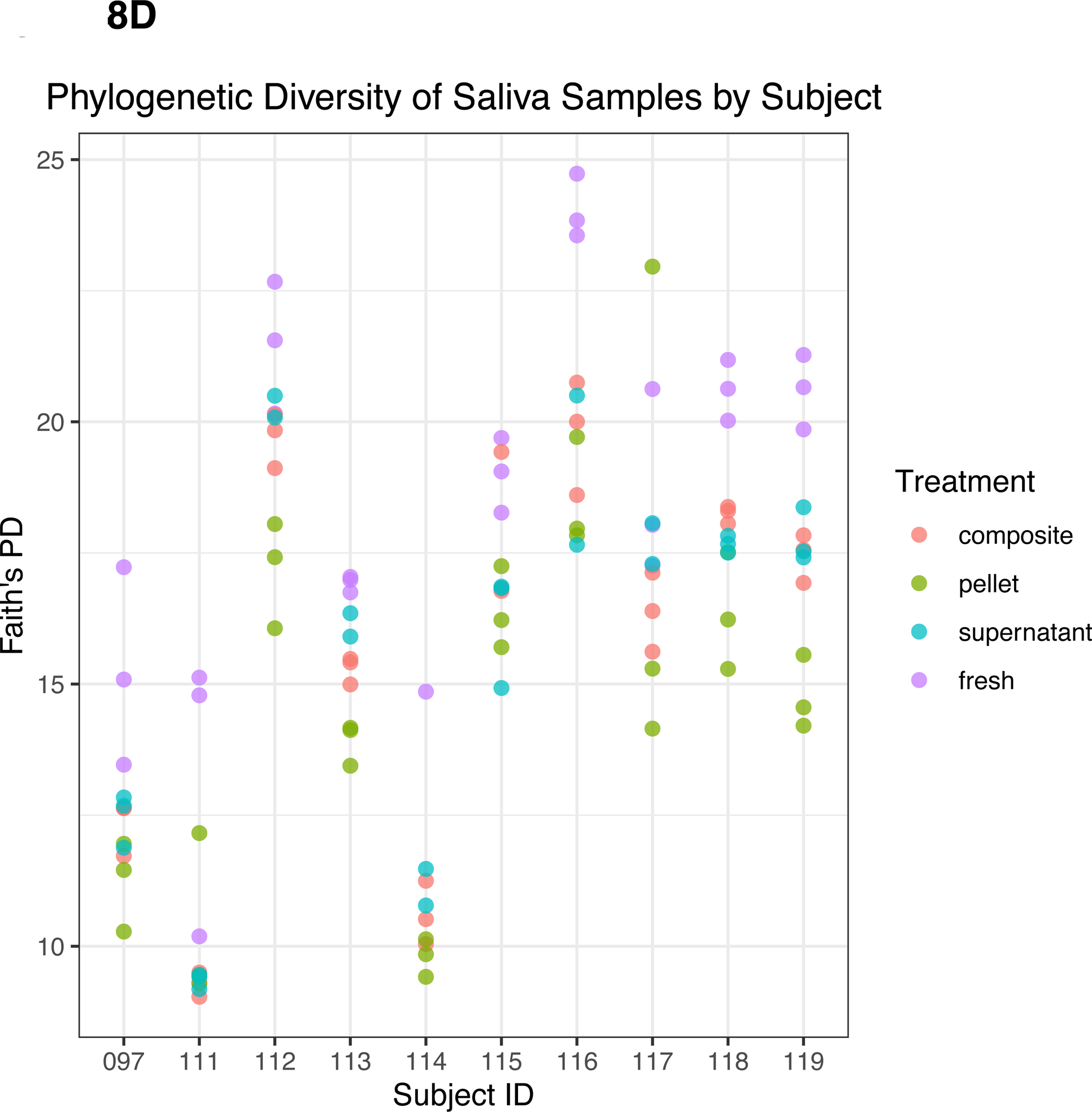
Alpha diversity of saliva samples. (A) Observed ASVs in saliva samples by treatment. (B) Evenness in saliva samples by treatment. (C) Phylogenetic diversity as determined by Faith’s PD in saliva samples by treatment. Statistically significant (p<0.05) differences are denoted by an asterisk (*) in A-C. (D) Phylogenetic diversity by individual subject for each saliva sample type.

### Differential abundance analysis of highest diversity samples

In addition to differences in diversity metrics, we identified microbes at the ASV level that distinguished the samples in the processing conditions with the highest alpha diversity from the others. Thus, for fecal samples, we compared fresh, LN2, and LN2post72hr samples to lyophilized. After FDR-adjusting p-values to correct for multiple comparisons, there were 230 ASVs with relative abundances that were significantly different from lyophilized in total (92 ASVs differed between fresh and lyophilized, 61 ASVs differed between LN2 and lyophilized, and 77 ASVs differed between LN2post72hr and lyophilized; adjusted p, q<0.05) (Supplementary Table 1). The ASV that was most increased in relative abundance in both fresh and LN2post72hr compared to lyophilized was assigned to *Prevotella copri*. The ASV that was most increased in relative abundance in LN2 compared to lyophilized was assigned to *Bacteroides plebeius*.

For saliva samples, we compared composite, pellet, and supernatant to fresh. There were 145 ASVs that significantly differed from fresh in total (54 differed between pellet and fresh, 43 differed between composite and fresh, and 48 differed between supernatant and fresh; q<0.05) (Supplementary Table 1). The two ASVs that were most increased in relative abundance in pellet and composite compared to fresh were both assigned to *Veillonella dispar*, and the ASV that was most increased in relative abundance in supernatant compared to fresh was assigned to *Selenomonas*. For saliva samples we also compared composite and pellet to supernatant, and we found 21 ASVs that were significantly different between composite and supernatant, and 70 ASVs were significantly different between pellet and supernatant (q<0.05). The same ASV that was most increased in relative abundance in both composite and pellet compared to supernatant was assigned to *Prevotella*.

For dental samples, we compared composite and pellet to supernatant, our reference condition, as it had the highest alpha diversity among the sample fractions. There were 3 ASVs that were significantly different between supernatant and composite, and 33 ASVs that were significantly different between supernatant and pellet, (q<0.05) (Supplementary Table 1). Out of these significant results, the ASV that was most increased in relative abundance in composite compared to supernatant was assigned to Pasteurellaceae. The ASV that was most increased in relative abundance in pellet compared to supernatant was an ASV assigned to *Actinobacillus* (in the Pasteurellaceae family). All statistically significant results are presented in (Supplementary Table 1).

A summary of our above findings can be found in Table 1.

**Table 1.** Summary of findings

| Sample type | Treatment or fraction with highest phylogenetic diversity | Significantly different ASVs |
| --- | --- | --- |
| Fecal | Lyophilized | 230 ASVs different between lyophilized and the other three treatments (fresh, LN2, |
|  |  | and LN2post72hr)<br>• 92 ASVs different between fresh and lyophilized<br>• 61 ASVs different between LN2 and lyophilized<br>• 77 ASVs different between LN2post72hr and lyophilized |
| Dental | Supernatant | 36 ASVs different between supernatant and the other two treatments (composite and pellet)<br>• 3 ASVs different between composite and supernatant<br>• 33 ASVs different between pellet and supernatant |
| Saliva | Fresh | 145 ASVs different between fresh and the other three treatments (pellet, composite, and supernatant)<br>• 54 ASVs different between pellet and fresh<br>• 43 ASVs different between composite and fresh<br>• 48 ASVs different between supernatant and fresh |

### Correlations between saliva and gut ASVs

Finally, we performed a correlation analysis on ASVs from saliva and fecal samples, due to a previous report of the impact of salivary amylase on both ends of the digestive tract (Poole et al., 2019), as well previous studies that indicate oral and gut microbiomes can be correlated in certain conditions such as type 1 diabetes (Kunath et al., 2022). For this analysis we compared the highest diversity saliva sample type with the highest diversity fecal sample type. After filtering ASVs to only include those that were present in greater than 50% of participants, we compared the 116 ASVs present in fresh saliva samples and 169 ASVs present in lyophilized fecal samples and found that out of the 19,604 comparisons, 50 ASVs were correlated with an FDR-adjusted p<0.05. All statistically significant results are shown in Supplementary Table 2.

## DISCUSSION

Our findings indicate that comparing microbial composition results across studies should be done with care when the storage or processing methods differ. We found that treatment methods, including both storage and processing methods, significantly affected diversity metrics and the differential abundance of ASVs between treatments in dental swabs, saliva, and fecal samples. Our most notable result was that lyophilized samples had microbiomes that were most distinct from and had the highest phylogenetic diversity and evenness compared to the same samples that underwent the other treatment methods (with the exception of Faith’s PD between lyophilized and LN2post72hr). Based on our results, lyophilization may be the preferred method for maximizing the diversity observed in fecal samples. However, caution should be exercised because not only does lyophilization result in higher alpha diversity than the other methods (likely due to more efficient lysis of bacterial cells), but it does so in a preferential manner, as lyophilized fecal samples had a significantly greater proportion of Firmicutes to Bacteroidetes than samples in all three other treatment methods. Although the Firmicutes to Bacteroidetes ratio has been posited as a marker of obesity (Ley et al., 2005; Mathur and Barlow, 2015; Koliada et al., 2017), this association is not consistent (Finucane et al., 2014; Sze and Schloss, 2016; Johnson et al., 2017; Magne et al., 2020). Our observation that a technical factor can affect this ratio further supports caution in the use of the Firmicutes to Bacteroidetes ratio as a biomarker.

Lyophilized fecal samples also had a significantly greater relative abundance of Archaea compared to samples in the other treatments, therefore indicating the ability of this treatment to increase the detection of microbes from this domain. Of note, having higher amounts of methanogenic Archaea in the gut has been correlated with obesity (Zhang et al., 2009; Basseri et al., 2012; Mathur and Barlow, 2015). It has also been reportedly unclear as to whether the predominant archaeal species *Methanobrevibacter smithii* is only present in a certain percentage of the population or whether a technical factor has hindered its detection (Gaci et al., 2014).

We also found that lyophilized fecal samples had less *Prevotella copri* compared to fresh and LN2post72hr samples. Previous studies suggest *P. copri* as an important marker of various health-related conditions and outcomes. In Kovatcheva-Datchary et al., individuals with gut microbiomes enriched with *P. copri* had improved glucose metabolism following the consumption of dietary fiber-enriched bread, and gavaging mice on a high fiber diet with *P. copri* resulted in improved glucose tolerance (Kovatcheva-Datchary et al., 2015). Another study showed that mice treated with *P. copri* resulted in lower glucose levels compared to the control group (Verbrugghe et al., 2021). Increased *P. copri* has also been suggested to play a role in rheumatoid arthritis (Scher et al., 2013; Jiang et al., 2022). In light of these examples, it is imperative to take fecal sample treatment type into account when comparing results between studies. Finally, for fecal samples, our beta diversity analysis revealed that treatment condition better explains the variation between samples when microbial abundance is taken into consideration (weighted UniFrac) rather than just presence or absence, which suggests that the effect of treatment is more prominent when giving less weight to low abundance ASVs.

Although distinction between donors was generally maintained between treatments for oral samples, we detected some ASVs that were significantly enriched depending on the treatment. Furthermore, supernatant had the highest alpha diversity out of the three frozen fractions for both saliva and dental samples, and therefore it may be beneficial to use this fraction over the others if presented with the option to do so. *Selenomonas* was found to be the ASV that was most increased in relative abundance in thawed saliva supernatant compared to fresh. This genus includes species such as *Selenomonas noxia*, which has been associated with periodontitis disease (Tanner et al., 1998; Torresyap et al., 2003; Cruz et al., 2015). Fresh saliva had higher alpha diversity than any of the frozen fractions. However, we recommend using thawed supernatant if using fresh samples that have never been frozen is impossible. We acknowledge that it is rarely feasible, especially for high throughput studies, to conduct DNA extractions immediately after receipt of a sample in the lab, particularly as samples are often not collected all at once. It is important to note that in this study the centrifugation speed that separated the supernatant and the pellet was relatively low (1500 x g at 4°C for 15 minutes), and some protocols might recommend much higher centrifugation speeds to form a pellet in order to maximize DNA yield, in which case the supernatant may not contain as much DNA.

There is no “one size fits all” recommendation. The detection of more distinct microbes in samples could be indicative of an observed microbiome that is closest to the true composition of the sample *in vivo*; therefore, investigators may wish to choose methods that maximize alpha diversity. However, in some cases, such as with lyophilization of fecal samples, the compositional differences from other methods could be a drawback when lyophilized samples are being analyzed with samples processed using other methods in a meta-analysis.

In conclusion, multiple different metrics of microbiome composition were affected by treatment methods for both gut and oral microbiome samples, with some methods increasing observed microbial diversity over others and enriching a subset of ASVs that have been associated with various aspects of host health. Therefore, it is important to exercise caution when selecting a processing method for one’s own study as well as when comparing microbial composition results across studies that use different processing methods.

## AUTHOR CONTRIBUTIONS

ACP designed the study. ACP and WZ performed experiments. DKS analyzed the data and performed the statistical analysis. DKS and ACP wrote the manuscript. All authors reviewed and approved the final manuscript prior to submission.

## ACKNOWLEDGEMENTS

We thank the Genomics Facility (RRID:SCR_021727) of the Biotechnology Resource Center of Cornell Institute of Biotechnology for performing sequencing and the Cornell Statistical Consulting Unit for their statistics expertise.

